# Dysregulated gene expression through *TP53* promoter swapping in osteosarcoma

**DOI:** 10.1101/2020.04.20.050252

**Authors:** Karim H. Saba, Valeria Difilippo, Michal Kovac, Louise Cornmark, Linda Magnusson, Jenny Nilsson, Hilda van den Bos, Diana C. J. Spierings, Mahtab Bidgoli, Tord Jonson, Vaiyapuri P. Sumathi, Otte Brosjö, Johan Staaf, Floris Foijer, Emelie Styring, Michaela Nathrath, Daniel Baumhoer, Karolin H. Nord

**Author notes:** These authors contributed equally to this work (Valeria Difilippo, Michal Kovac). D.B. and K.H.N. jointly directed this work.

## Abstract

How massive genome rearrangements confer a competitive advantage to a cancer cell has remained an enigma. The malignant bone tumour osteosarcoma harbours an extreme number of structural variations and thereby holds the key to understand complex cancer genomes. Genome integrity in osteosarcoma is generally lost together with disruption of normal *TP53* gene function, the latter commonly through either missense mutations or structural alterations that separate the promoter region from the coding parts of the gene. To unravel the consequences of a *TP53* promoter relocated in this manner, we performed in-depth genetic analyses of osteosarcoma biopsies (*n*=148) and cell models. We show that *TP53* structural variations are early events that not only facilitate further chromosomal alterations, but also allow the *TP53* promoter to upregulate genes erroneously placed under its control. Paradoxically, many of the induced genes are part of the *TP53*-associated transcriptome, suggesting a need to counterbalance loss of *TP53* function through ‘separation-of-function’ mutations via promoter swapping. Our findings demonstrate how massive genome errors can functionally turn the promoter region of a tumour suppressor gene into a constitutively active oncogenic driver.

## Introduction

Osteosarcoma is the most common primary malignancy of the skeleton. The majority of osteosarcomas develop in children and adolescents, often in close proximity to the active growth plate of long bones^1^. In the third decade of life, the incidence rate drops but is followed by a second, smaller incidence peak in elderly individuals. During the 1980s, the introduction of multidrug chemotherapy dramatically improved the survival rate of osteosarcoma patients. Clinical outcome has improved little since then, however. The overall survival rate remains at 60-80% for localised disease and below 40% for disseminated osteosarcoma^1^. Osteosarcoma typically displays a very large number of numerical and structural chromosome aberrations, often in the several hundreds across the genome^2–8^. Hitherto, there are no reports on genetic alterations specific to this disease and a consistent genetic pattern between patients is lacking, something that has hampered the identification of novel therapeutic targets.

Large-scale sequencing efforts have reliably demonstrated that the vast majority of osteosarcomas harbour mutations in the *TP53* gene^3, 5^, the master guardian of genome integrity. Some of these are hotspot mutations that encode missense mutant p53 proteins. Although the cause and pathogenetic significance of such mutations are not fully understood, they are common features of many high-grade cancers. Emerging evidence suggest that they could constitute gain-of-function or separation-of-function mutations^9–13^. The latter type of mutation implies that while the classical tumor supressor actitivites of *TP53* are lost, survival and proliferative features of the *TP53* pathway are retained. Paediatric osteosarcomas are unique among high-grade malignancies in that at least half of the cases show structural variations in *TP53*^2, 3, 5^. These rearrangements separate the promoter region from the coding parts of *TP53*, often resulting in loss of the latter. Interestingly, the promoter region is not lost, but instead relocated, thereby enabling the erroneous activation of genes other than those originally under its control. Such transfer of promoter activity is a known driver in neoplasia, commonly denoted promoter swapping/switching or enhancer hijacking. Promoter swapping has been shown to operate in bone tumours other than osteosarcoma, *e.g.*, in chondromyxoid fibroma and aneurysmal bone cyst where strong promoters are juxtaposed to the entire coding sequences of the *GRM1* and *USP6* genes, respectively^14, 15^. Previously reported promoter substitutions in neoplasia have typically involved supposedly ‘strong’ promoters, assumed to be constitutively active in the cell-of-origin^16^. Here, we use the complex genome of osteosarcoma to test the novel hypothesis that acquired genetic damage can activate a transferred promoter of a tumour suppressor to drive oncogenesis.

## Results

### Ectopic localisation of the *TP53* promoter is prevalent in young osteosarcoma patients

To make a detailed assessment of the role of *TP53* rearrangements in osteosarcoma, we first subjected a discovery cohort of conventional osteosarcomas from paediatric (age <18 y, *n*=15) and adult (age range 18-81 y, *n*=21) patients to whole genome mate pair sequencing, which is a powerful technology to identify structural genomic alterations. The majority of samples analysed in this cohort were chemotherapy-treated resection specimens. We found structural rearrangement of *TP53* in 13/36 cases (Fig. 1a; Discovery cohort, Supplementary Tables 1 and 2). We then analysed an independent validation cohort of treatment-naïve diagnostic biopsies from conventional osteosarcomas, again including both paediatric (age <18 y, *n*=20) and adult (age range 18-59 y, *n*=16) patients. In the validation cohort, structural rearrangement of *TP53* was found in 16/36 cases (Fig. 1a; Validation cohort, Supplementary Tables 3 and 4). We then extended our validation cohort and analysed genome-wide DNA copy number profiles based on SNP array data from these cases (age range 3-74 y, *n*=108; Supplementary Table 3). By integrating array and sequencing data, we identified a subset of cases with a distinct copy number profile of chromosome arm 17p that we termed *‘TP53* promoter gain’. We defined this pattern as copy number loss, or copy number neutral loss of heterozygosity, of whole or parts of the *TP53* coding region coupled to concurrent relative copy number gain of the *TP53* promoter region along with regions of proximal chromosome arm 17p (Fig. 1b). We found *TP53* promoter gain in 16/108 cases (15%; Fig. 1c). Both *TP53* promoter gain, determined by SNP array analysis, and *TP53* structural variation, determined by whole genome mate pair sequencing, were non-randomly associated with young age of onset (Fig. 1d-e, Supplementary Tables 1 and 3). In an additional 24 of the 108 tumours analysed by SNP array (22%), we detected a copy number shift within the nearest measuring points downstream and upstream relative to *TP53* but lacking at least one criterion for *TP53* promoter gain (Supplementary Table 3). Collectively, we identified *TP53* structural variants in 40% of conventional osteosarcomas, *i.e.*, 29/72 by DNA mate pair sequencing and 40/108 by SNP array analysis (Supplementary Tables 1 and 3). Additionally, a breakpoint burden analysis showed that cases with a *TP53* structural variant had a higher number of chromosome breaks genome-wide than cases without one (Fig. 1f, Supplementary Tables 1 and 3). A more in-depth analysis of breakpoint distributions across the genomes of the 72 sequenced osteosarcomas revealed multiple regions with a marked clustering of breakpoints, including chromosome arms 6p, 8q, 12q, 17p and 19q (Supplementary Fig. 1). There was a sharp contrast between regions proximal and distal to *TP53*, where the number of breakpoints increased substantially from *TP53* intron 1 and towards the centromere (Fig. 1g), closely mimicking the genomic copy number pattern illustrated in Fig. 1c. A representative case is depicted in Fig. 1h.

**Figure 1.**
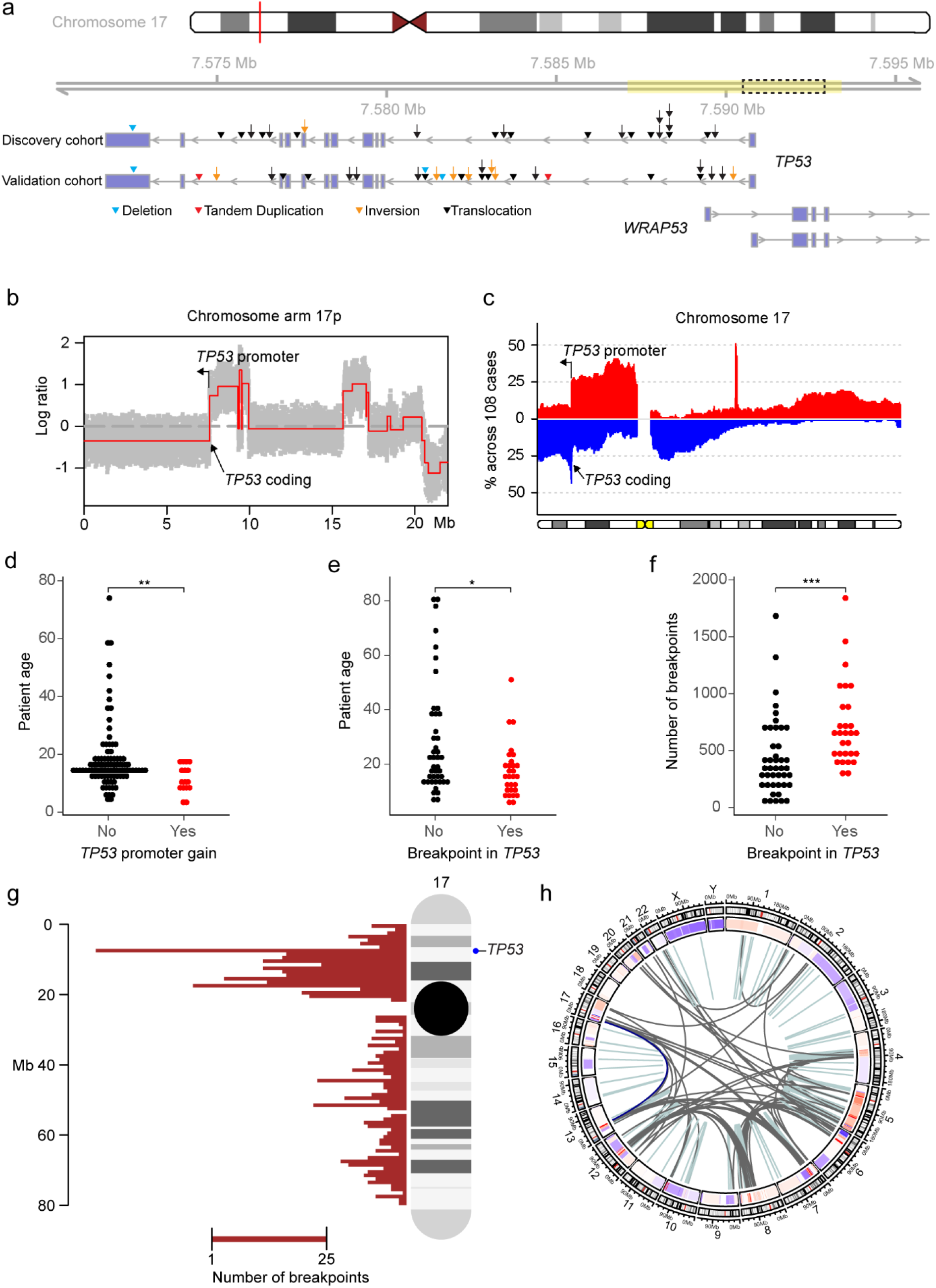
Structural variation in *TP53* is associated with young age at onset and a high number of chromosomal breaks genome-wide. **a** Schematic representation of *TP53* structural variation in a discovery (*n*=36) and a validation osteosarcoma cohort (*n*=36). The *TP53* promoter region is marked in yellow^67^. The region used to represent the *TP53* promoter region in *in vitro* experiments is marked by a dashed box. Arrowheads and arrows represent structural variants involving sequences 5’ and 3’ of the breakpoint, respectively. **b** DNA copy number profile of 17p in a representative osteosarcoma with gain of the *TP53* promoter region. **c** Frequency plot of genomic copy number gain (red) and loss (blue) for chromosome 17 across conventional osteosarcomas (*n*=108). **d** Age distribution of osteosarcoma patients without (*n*=92) and with (*n*=16) *TP53* promoter gain as determined by SNP array analysis. \*\**P* < 0.01, two-tailed Mann-Whitney *U* test. **e** Age distribution of osteosarcoma patients without (*n*=43) and with (*n*=29) *TP53* structural variants as determined by DNA mate pair sequencing. \**P* < 0.05, two-tailed Mann-Whitney *U* test. **f** Breakpoint burden distribution of osteosarcomas without (*n*=43) and with (*n*=29) *TP53* structural variants as determined by DNA mate pair sequencing. \*\*\**P* < 0.001, two-tailed Mann-Whitney *U* test. **g** All detected breakpoints affecting chromosome 17 across conventional osteosarcomas (*n=*72). Red histograms represent total counts within a specific genomic window. **h** Circos plot showing genome rearrangements in a representative osteosarcoma with structural variation in *TP53*. Light blue and dark grey lines denote intra- and interchromosomal events, respectively. The dark blue line represents the specific structural variant relocating the *TP53* promoter region.

Cases that lacked a structural variant directly affecting the *TP53* locus displayed a diploid copy number of the gene, larger copy number losses or gains of chromosome arm 17p that did not directly affect *TP53* integrity, a complete homozygous loss of the gene, and/or single nucleotide variants or indels leading to missense or frameshift mutations (Supplementary Tables 1 and 3). *TP53* missense mutations were mutually exclusive from *TP53* structural variants.

The recurrent transposition of the *TP53* promoter suggested that it regulates other genes in a fashion that favours tumour development, through gene fusion or promoter swapping events^16^. To test this hypothesis, we performed several analyses. We assessed gene expression levels by RNA sequencing of conventional osteosarcomas from which good quality RNA could be obtained, including cases from the discovery and validation cohorts as well as a third cohort described below (age range 4-81 y, *n*=68; Supplementary Tables 1, 3 and 5). To evaluate the presence of *TP53* promoter gain in the discovery cohort, we analysed DNA copy numbers in cases from which material was available (*n*=34; Supplementary Table 1). To determine if *TP53* structural variations were present among multiple samples from the same tumour, we analysed a third cohort of five *TP53*-rearranged osteosarcomas sampled across several regions and time points. In these cases, we compared paired-end whole genome and RNA sequencing data from diagnostic biopsies, resection specimens, and/or metastases (*n*=11 biopsies; Supplementary Tables 5 and 6). To evaluate the proportion of *TP53* structural variations among individual cells from the same tumour, we finally applied single cell low-pass whole genome sequencing to cryopreserved cells from two osteosarcomas of the discovery cohort. By integrating the obtained high-resolution genomic data with matched transcriptomic information, we found that transposition of the *TP53* promoter is an early event that results in deregulation of several well-known or putative oncogenes of which some, paradoxically, are part of the *TP53*-signalling pathway. Below we provide several lines of evidence for this mechanism.

### Transposition of the *TP53* promoter is a single early event that can spark genome-wide rearrangements and oncogene amplification

In a subset of osteosarcomas, DNA sequencing supported intra- and interchromosomal events (inversions, insertions or translocations) that transposed the *TP53* promoter without compromising chromosome stability (Fig. 2a-c). In these cases, we detected no further rearrangements involving the *TP53* promoter or its partner region. In another subset of osteosarcomas, transposition of the *TP53* promoter was the initiating event that generated unstable, most likely dicentric, derivative chromosomes (Fig. 2d-f, Supplementary Fig. 2 and 3). In osteosarcoma, such derivative chromosomes repeatedly break and re-join with multiple partner chromosomes^17, 18^. This amplifies both the *TP53* fusion and additional genomic regions of potential importance for osteosarcoma progression, such as regions on chromosomes 6, 12 and 17 (Fig. 2f). Notably, this sequence of events is different from chromothripsis and multi-way translocations, which in other subtypes of bone tumours are known to generate gene fusions (Fig. 2g-h) ^19, 20^. We found no evidence for the generation of *TP53* structural variants or *TP53* gene fusions through one massive burst of genome rearrangements in osteosarcoma. Instead, the genomic footprint of *TP53* gene fusions in osteosarcoma mimics that of oncogene amplification through breakage-fusion-bridge cycles, found in *e.g.*, low-grade osteosarcoma with ring chromosomes and *MDM2* amplification (Fig. 2i). Thus, according to our model, transposition of the *TP53* promoter is an early spark for genome-wide rearrangements in osteosarcoma. Results from whole genome sequencing of multi-sampled bulk and single cell tumour DNA supported this model. *TP53* fusion positive osteosarcomas harboured their respective fusions in all investigated diagnostic biopsies, post-chemotherapy resection specimens and metastases, as well as in all investigated individual neoplastic cells (Fig. 2j-k, Supplementary Fig. 3a-f, 4-6 and Supplementary Tables 3 and 5).

**Figure 2.**
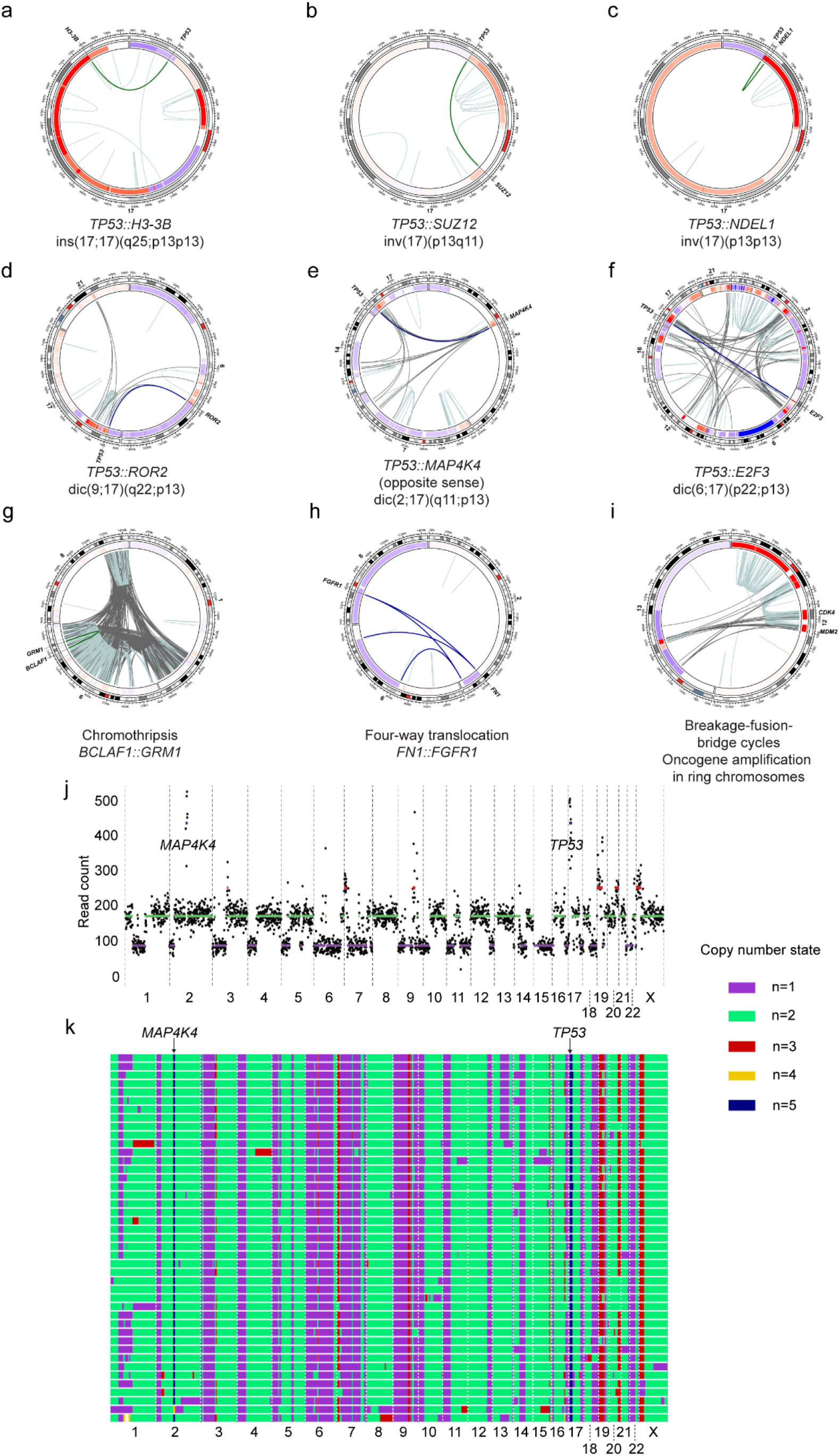
Transposition of the *TP53* promoter is a single early event that can spark genome-wide rearrangements and oncogene amplification. **a-c** Intrachromosomal events resulting in *TP53* gene fusions (green lines). **d-f** Interchromosomal events resulting in *TP53* gene fusions (dark blue lines). The derivative dicentric chromosomes repeatedly break and re-join with multiple partner chromosomes. Exemplified are the genomic footprints of **(g)** chromothripsis in a chondromyxoid fibroma, **(h)** a multi-way translocation in a phosphaturic mesenchymal tumour of bone and **(i)** breakage-fusion-bridge cycles in a parosteal osteosarcoma. **j** Genomic copy numbers in a representative individual cell from an osteosarcoma with a *TP53::MAP4K4* fusion. **k** Heat map of genomic copy numbers across all 43 sequenced individual neoplastic cells of the *TP53::MAP4K4* fusion positive case. Each row of copy number states represents a single cell.

### The bidirectional *TP53* promoter induces the expression of *WRAP53* and partner genes *in vivo*

To assess if the *TP53* promoter induces the expression of its respective partner genes *in vivo*, we analysed the expression levels for the partner genes and the *WRAP53* gene. The *TP53* promoter is bidirectional and normally induces both *TP53* and *WRAP53*^21^, wherefore elevated expression levels of the latter were used as a proxy for adequate representation of neoplastic cells. Fig. 3a-f displays three representative osteosarcomas that harbour whole or parts of *ROR2, MAP4K4* and *E2F3*, respectively, placed under the control of the *TP53* promoter. *TP53* exon 1 and partner gene exons placed under the *TP53* promoter showed higher expression levels than exons excluded from the fusions. Other notable genes that were placed under the control of the *TP53* promoter included *e.g.*, *ELF1, CDC5L, H3-3B, YTHDF1, ZNF780A, SUZ12, NDEL1, CDKN1A* and *NFYA.* Although none of the 3’ partners were recurrent in themselves, their respective functions seemed to merge on common pathways. For example, several partner genes have been shown to regulate cartilage and growth plate development, osteoclast formation or to be implicated in cancer development in other tumour types^22–31^. Furthermore, and perhaps even more intriguingly, pathway enrichment analyses indicated that most *TP53* promoter partner genes in cases with *TP53* promoter gain are themselves part of the *TP53*-signalling pathway (Fig. 3g and Supplementary Table 7). In line with this, the global gene expression pattern of individual osteosarcomas did not correlate with their *TP53* mutation status (Supplementary Fig. 7). Thus, in *TP53*-mutated cases, be it by structural or single nucleotide variants, a compensatory mechanism could potentially restore, at least partly, the *TP53* wildtype phenotype. In the supplementary Tables 1, 3 and 5, we display the matched genomic and transcriptomic data for all detected *TP53* gene fusions. A schematic showing which genes (or parts of genes) are placed under the control of the *TP53* promoter in all cases harbouring a *TP53* structural variant is provided in the supplementary Fig. 8. Taken together, these data unequivocally demonstrate that the transposed *TP53* promoter is active in osteosarcoma and that it induces the expression of genes important for tumour and bone development which, in some instances, are themselves involved in the *TP53*-signalling pathway.

**Figure 3.**
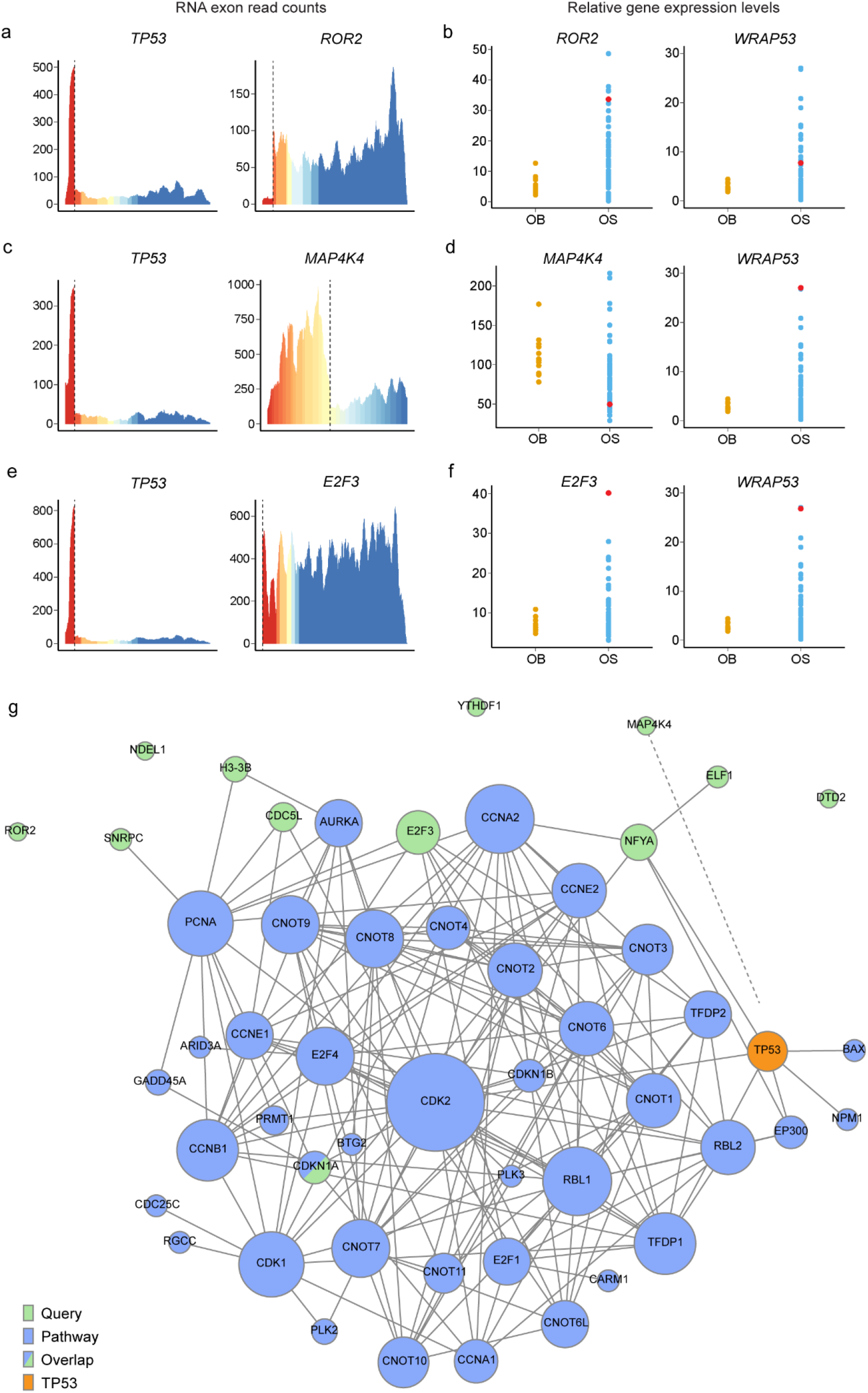
The bidirectional *TP53* promoter induces the expression of *WRAP53* and oncogenes *in vivo.* **a** Exon expression levels in Case 9, in which *TP53* intron 1 is fused to *ROR2* exons 2-9. **b** Normalised gene expression levels. **c** Exon expression levels in Case 22, in which *TP53* intron 1 is fused to *MAP4K4* exons 1-15, including coding regions for the kinase domain, in the opposite direction. **d** Normalised gene expression levels, including all exons of *MAP4K4* in Case 22. **e** Exon expression levels in Case OS046, in which *TP53* intron 1 is fused to regions upstream the complete coding sequence of *E2F3.* **f** Normalised gene expression levels. Different colours mark individual exons. Dotted lines indicate the fusion points. Red dots mark the case under investigation. OB = osteoblastoma, OS = osteosarcoma. **g** The *TP53* promoter partners *ELF1, NFYA, E2F3, CDKN1A, CDC5L, H3-3B, SNRPC* and *MAP4K4* are connected to the *TP53*-signalling pathway, either as direct downstream effectors of TP53 or several steps downstream. Genes marked in blue are part of the *TP53* pathway while those marked in green are the *TP53* promoter partners that were queried against the MSigDB and PathBIX database. Overlaps are marked in both blue and green. P < 0.001, FDR < 0.05.

### Cisplatin evokes gene expression through the *TP53* promoter *in vitro*

As a proof-of-concept, we modelled the above findings *in vitro.* First, we knocked out *TP53* in one mesenchymal (BJ-5ta) and one epithelial (RPE-1) cell line by CRISPR genome editing and single cell cloning. Second, we constructed a vector containing the *TP53* promoter region fused to the coding DNA sequence of *ROR2 (TP53::ROR2).* As a control for the absence of the promoter, we used the same vector but without the *TP53* promoter region (*ROR2*). Third, we exposed *TP53^-/-^* cells harbouring either *TP53::ROR2* or *ROR2* to the DNA damaging agent cisplatin (Fig. 4a-b). We found that the *TP53^-/-^* background, even in the absence of cisplatin, was sufficient to activate the *TP53* promoter and elicit expression of a gene placed under its control (Fig. 4c-d). Induced DNA damage through cisplatin treatment further increased the expression level of the *TP53* promoter partner gene. The same phenomenon was detected in the osteosarcoma cell line Saos-2, known to harbour the *TP53::SAT2* fusion^5^ (Fig. 4e). In the highly proliferative osteosarcoma cell line MG-63 harbouring a *TP53::ADGRE1* (formerly *EMR1*) fusion^5, 32^, there was a clear upregulation of the partner gene compared to control (Fig. 4f). However, these cells were heavily affected by the cisplatin treatment and the *ADGRE1* expression level decreased with increasing cisplatin concentration. A probable explanation for this effect could be that their high proliferation rate renders them more susceptible to cisplatin. Thus, in a *TP53^-/-^* background, a constitutively active *TP53* promoter can induce expression of an oncogene transposed into its vicinity in a fashion that can be accentuated by additional genetic damage.

**Figure 4.**
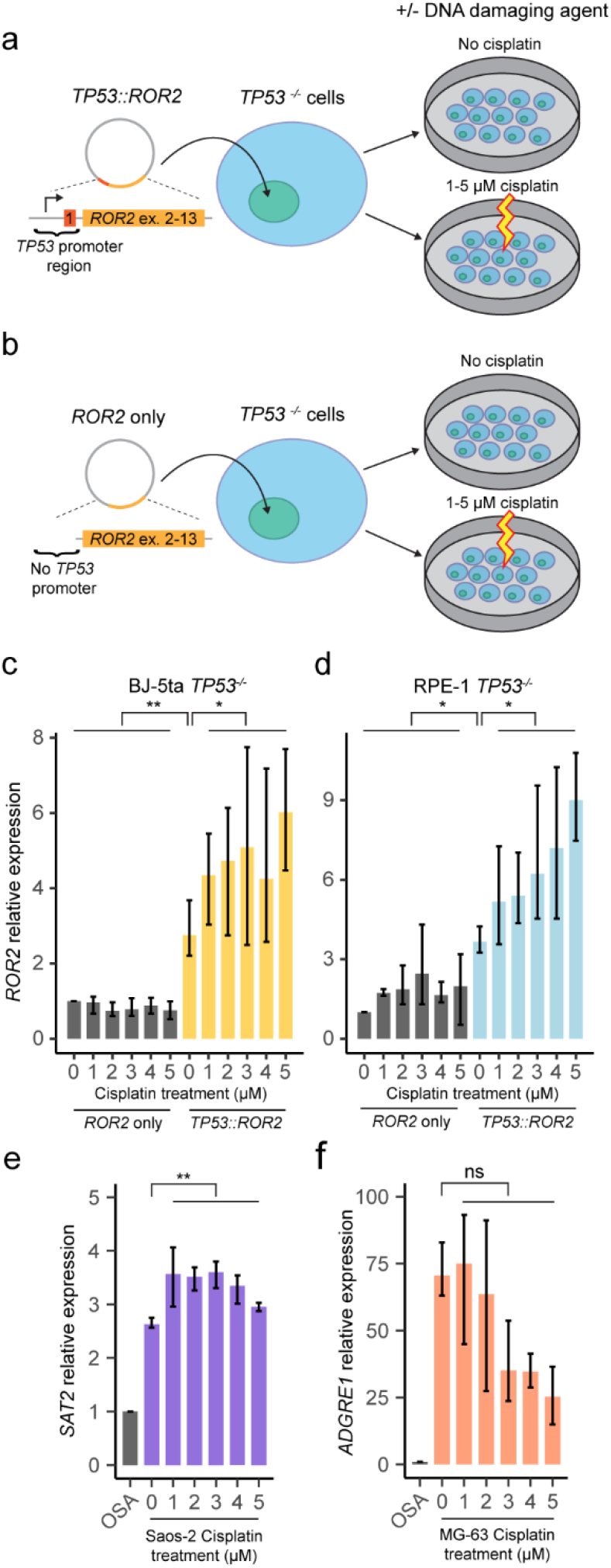
A *TP53* null background constitutively activates the *TP53* promoter. **a-b** Bj-5ta *TP53^-/-^* and RPE-1 *TP53^-/-^* cells were lentivirally transduced with a promotor-less vector containing either (a) the *TP53* promoter region fused to *ROR2* exons 2-13, or (b) *ROR2* exons 2-13 without the *TP53* promoter region. Cells were then cultured with or without 1-5 μM cisplatin for four days, followed by RNA extraction. RT-qPCR was used to measure the **c** *ROR2* relative expression levels in BJ-5ta *TP53^-/-^* cells, **d** *ROR2* relative expression levels in RPE-1 *TP53^-/-^* cells, **e** *SAT2* relative expression levels in Saos-2, and f *ADGRE1* relative expression levels in MG-63. *n* = 3 biological replicates, mean ± range, \**P* < 0.05, \*\**P* < 0.01, ns = nonsignificant, two-tailed Mann-Whitney *U* test.

### Introduction of *TP53::ROR2* in *TP53* null cells reverts their global gene expression profile back towards the wildtype phenotype

Finally, we used whole transcriptome sequencing to assess the global gene expression patterns in cultured wildtype Bj-5ta, BJ-5ta *TP53^-/-^*, and BJ-5ta *TP53^-/-^* harbouring *TP53::ROR2* cells. Unsupervised principal component analysis showed that knockout of *TP53* clearly affected the global gene expression profile, i.e., there was a clear distinction between the wildtype cells and those with a complete knockout of *TP53* (Supplementary Fig. 9). Intriguingly, in *TP53^-/-^* cells that harboured the *TP53::ROR2* fusion, the global gene expression profile reverted back towards that of wildtype cells. This is in line with the above findings that *TP53* promoter swapping events potentially counterbalance loss of *TP53* function through separation-of-function mutations.

## Discussion

The first reports on *TP53* structural rearrangements in osteosarcoma date back to the late 1980s and early 1990s^33, 34^. Already then, the clustering of alterations to *TP53* intron 1 was noted and it was speculated that ‘rearrangements of p53 in osteosarcoma could activate a second as yet unidentified gene’^34^. During the following decades, efforts from several research groups confirmed these rearrangements, and genomic patterns similar to what we here term ‘*TP53* promoter gain’ were reported in osteosarcoma and subtypes of soft tissue sarcomas^35^. In parallel, somatic structural variations affecting *TP53* were also found in subsets of leukaemia and carcinomas, including chronic myelogenous leukaemia^36–38^, lung cancer^39^ and prostate cancer^40–43^. Such variants inevitably silence the *TP53* gene, but evidence for concomitant gain-of-function or separation-of-function mechanisms through structural variation has not been described. This is likely because essentially all studies investigating *TP53* gain- or separation-of-function mutations were focused on single nucleotide variants^9–11^. Here, we propose that dislocating the *TP53* promoter region from the coding parts of the gene represents an effective means to accomplish separation-of-function mutations, particularly in young osteosarcoma patients who have not lived the long life thought necessary to accumulate a high number of somatic single nucleotide variants. By distorting normal *TP53* function while simultaneously inducing cancer-related signalling pathways, structural variations of the *TP53* region likely mirror the mode of action of many *TP53* missense mutants^11^. For *TP53* structural variants and missense mutations to come into play as gain-of-function or separation-of-function mutations, the cancer cell must likely be deprived of the production of normal *TP53* through losses, nonsense or frameshift mutations affecting the other allele.

There may be at least two probable reasons for not recognizing *TP53* separation-of-function mutations through structural variation in previous studies. First, the *TP53* promoter is a promiscuous fusion partner that induces the expression of many different genes. This, however, does not exclude an important functional outcome. There are numerous examples of interchangeable partners of gene fusions that are disease-specific, strongly indicating that activation of a specific pathway, in one way or the other, is the key feature for transformation^16, 44^. Second, the *TP53* gene fusions in osteosarcoma involve transfer of promoter activity. Although a well-recognised concept in neoplasia, its detection requires access to matched high quality genomic and transcriptomic data. We generated a unique combined dataset for a large series of paediatric and adult osteosarcomas, sampled across several regions and time points. This enabled us to show for the first time that a promoter activated by genetic damage can induce cancer-driving genes transposed into its vicinity. Importantly, we found this phenomenon to occur in all neoplastic cells of *TP53*-rearranged osteosarcomas, rendering it a particularly meaningful mechanism to explore further for therapeutic applications.

## Methods

### Subject information and tumour material

Fresh-frozen tumour biopsies from 148 conventional osteosarcomas were subjected to genomic analyses. The clinical features were typical of conventional osteosarcoma patients. The age of the patients ranged from 3-81 years with a median age of 15 years and a mean age of 20 years, and there were 68 females and 80 males. Detailed information is displayed in Supplementary Tables S1, S3 and S5. For comparison, we included osteoblastomas (*n*=13), a chondromyxoid fibroma, a phosphaturic mesenchymal tumour of bone and a parosteal osteosarcoma. All tumour material was obtained after informed consent, and the study was approved by the Regional Ethics Committee of Lund University and the Ethikkommission beider Basel (reference 274/12).

### DNA and RNA extractions

Fresh-frozen tumour biopsies were dismembered and homogenised using a Mikro-Dismembrator S (Sartorius AG, Goettingen, Germany). The material was optimally split into two fractions, one used for immediate DNA extraction and the other, when available, was stored in Qiazol at −80°C for later RNA extraction (Qiagen, Hilden, Germany). DNA was extracted using the DNeasy Blood & Tissue Kit including the optional RNase A treatment (Qiagen). DNA quality and concentration were measured using a NanoDrop ND-1000 and a Qubit 3 Fluorometer (Thermo Fisher Scientific, Waltham, MA). The material stored in Qiazol was heated at 65°C for 5 min and RNA was extracted using the RNeasy Lipid Tissue Kit including the optional DNase digestion (Qiagen). RNA quality and concentration were assessed using a 2100 Bioanalyzer (Agilent Technologies, Santa Clara, CA), and a NanoDrop ND-1000 (Thermo Fisher Scientific).

### Whole genome mate pair sequencing for detection of structural variations

To detect structural chromosomal abnormalities, mate pair libraries were prepared for sequencing using the Nextera Mate Pair Library Preparation Kit (Illumina, San Diego, CA). This was done according to the manufacturer’s instructions except for the number of shearing cycles, which were increased to three cycles. Paired-end 76 base pair reads were generated using an Illumina NextSeq 500 sequencing instrument. Sequencing depth was on average 3.2x (mapping coverage 2.13x) and the mean insert size was 3.0 kb, resulting in a median spanning coverage of 63.2x of the human genome (mean 63.1x, range 5.2x-119.1x). All samples were sequenced with high quality and yield; between 12.4 and 115.5 million read pairs were obtained per sample and the average quality scores were 31.3-34.1. Sequencing reads were trimmed using NxTrim v 0.4.2 and subsequently aligned against the GRCh37/hg19 build using the Borrows-Wheeler Aligner v 0.7.15^45^. To identify structural rearrangements, the sequence data were analysed using Integrative Genomics Viewer^46, 47^, as well as the structural variant callers TIDDIT v 2.12.1, Delly2 v 0.7.8 and Manta v 1.2.2^48–50^. Structural alterations were considered true when identified by at least two of the three variant callers.

### Whole genome paired-end sequencing of multi-sampled osteosarcomas

Whole genome paired-end sequencing was performed using the Agilent SureSelect v3 library preparation kit (Agilent Technologies, Santa Clara, CA). Paired-end 150 base pair reads were generated using an Illumina HiSeq 2500 sequencing instrument. Sequencing depth was on average 13.4x (mapping coverage 14.1x) and the mean insert size was 0.34 kb, resulting in a median spanning coverage of 14.5x of the human genome (mean 14.3x, range 5.2x-40.9x). Sequencing reads were aligned against the GRCh37/hg19 build using the Borrows-Wheeler Aligner v 0.7.15. To identify structural rearrangements, the sequence data were analysed as described above. It is important to stress that whole genome paired-end sequencing is a less optimal technique to detect structural variations, compared with mate pair sequencing, and therefore requires a higher sequencing depth. The reason is the higher spanning coverage of the human genome obtained by mate pair sequencing, due to the analysed DNA fragments being approximately one order of a magnitude larger. In the present study, the median spanning coverage for mate pair data was 63.2x compared to 14.5x for paired-end data.

### Genome-wide DNA copy number and loss of heterozygosity analyses

SNP array analysis was used for combined DNA copy number and loss of heterozygosity investigation. DNA was extracted according to standard procedures from fresh frozen tumour biopsies and hybridised to CytoScan HD arrays, following protocols supplied by the manufacturer (Thermo Fisher Scientific). Somatic copy number alterations in a proportion of the cases were published by Smida *et al.* 2017^7^. Data analysis was performed using the Chromosome Analysis Suite v 4.1.0.90 (Thermo Fisher Scientific), detecting imbalances by visual inspection, and by segmenting log2 values using the R package ‘copynumber’, available via Bioconductor. The inbuilt pcf function was used with a strict gamma value of 100 to create copy number segments and the plotFreq function was used to create the frequency plot of losses and gains on chromosome 17. The threshold for gain was set as a log2 value of 0.2 and the threshold for loss as −0.2. SNP positions were based on the GRCh37/hg19 sequence assembly. *‘TP53* promoter gain’ is defined as copy number loss, or copy number neutral loss of heterozygosity, of whole or parts of the *TP53* coding region coupled to concurrent relative copy number gain of the *TP53* promoter region along with regions of the proximal part of chromosome arm 17p.

### Visualisation of structural and copy number variations using circos plots

Circos plots were generated using the R packages ‘Circlize’ or ‘RCircos’, by integrating genomic copy number data obtained from either SNP array analysis or whole genome sequencing and structural variant data based on whole genome sequencing and the TIDDIT algorithm described above. Copy number segments based on SNP array data were generated as described above. Copy number segments based on sequencing data were generated using CNVkit^51^.

### Whole genome low-pass sequencing of single cells

Whole genome sequencing of cryopreserved primary osteosarcoma cells was performed as described in detail previously^52^. In brief, library preparation was performed using a modified single cell whole genome sequencing protocol and 77 base pair single reads were generated using a NextSeq 500 sequencing instrument (Illumina). From each assessed tumour, 93 individual cells were sequenced at an average depth of 0.01x. Copy number analysis was performed using AneuFinder^53^, and bin positions were based on the GRCh38/hg38 sequence assembly.

### RNA sequencing for detection of gene fusions, expression levels, single nucleotide variants and indels

Total RNA was enriched for polyadenylated RNA using magnetic oligo(dT) beads. Enriched RNA was prepared for sequencing using the TruSeq RNA Sample Preparation Kit v2 according to the manufacturer’s protocol (Illumina). Paired-end 151 base pair reads were generated from the cDNA libraries using an Illumina NextSeq 500 instrument. Sequencing reads were aligned to the GRCh37/hg19 build using STAR v 2.5.2b^54^. For comparison of relative gene expression levels, data were normalised using Cufflinks with default settings^55^, and visualised using the Qlucore Omics Explorer version 3.5 or 3.8 (Qlucore AB, Lund, Sweden). As an outgroup in gene expression analysis, we included osteoblastomas (*n*=13) in addition to the 68 osteosarcomas. FusionCatcher v 1.0 and STAR-Fusion v 1.4.0 were used to identify candidate fusion transcripts from the sequence data^56, 57^. Single nucleotide variants and indels in the *TP53* gene were detected using VarScan v 2.4.1 ^58^ and Mutect v 1.1.7^59^. Constitutional variants were excluded based on information from the Genome Aggregation Database (gnomAD v2.1.1) ^60^. The detected variants were finally confirmed by manual inspection using the Integrative Genomics Viewer. Unsupervised correlation-based principal component analysis was performed on all 68 osteosarcomas using the Qlucore Omics Explorer.

### PCR and Sanger sequencing

Genomic PCR, RT-PCR and nested RT-PCR for detection of the *TP53::ROR2, TP53::SUZ12, TP53::NDEL1*, and *TP53::RTBDN* gene fusions were performed as previously described^61^. Amplified fragments were purified from an agarose gel and Sanger sequencing was performed by GATC Biotech (Eurofins Genomics, Ebersberg, Germany). BLASTN software (https://blast.ncbi.nlm.nih.gov/Blast.cgi) was used for the analysis of sequence data.

### *In silico* verification of identified fusion transcripts: NAFuse

In order to determine the genomic support for fusion transcripts identified by RNA sequencing, we developed a pipeline which we named NAFuse. NAFuse is an integrated methodology that combines and compares the output of a gene fusion detector with that of the BAM-file generated by whole genome sequencing. This allows for automated detection of breakpoints in both partner genes/regions on both the RNA and DNA levels. The source code for NAFuse is available via GitHub. A detailed description of NAFuse and the link to the GitHub page can be found in the supporting information (supplementary materials).

### Pathway enrichment analyses

Genes identified as being partners to the *TP53* promoter region in cases with *TP53* promoter gain were queried as a list against the (i) Pathway Analysis with ANUBIX (PathBIX) database^62^, using the Reactome pathway database with a network cut-off set at 0.99 without applied clustering, and the (ii) Molecular Signatures Database (MSigDB) ^63–65^, where the H and C1-6 gene sets were used to compute gene overlaps.

### Cell models harbouring *TP53::ROR2, TP53::SAT2* or *TP53::ADGRE1*

A promoter-less vector (pSMPUW Universal Lentiviral Expression Vector, Cell Biolabs, Inc., San Diego, CA) containing the *TP53::ROR2* fusion was constructed (GenScript, Piscataway, NJ). The *TP53* promoter was represented by the first 2000 bp upstream of *TP53* together with exon 1 and the first 500 bp of intron 1 of *TP53.* These *TP53* sequences were fused to the last 500 bp of *ROR2* intron 1 and the coding sequences of *ROR2* exons 2-13. This hybrid sequence is denoted *TP53::ROR2* and thus contains the complete coding sequence of *ROR2* transcript variant 002 (ENST00000375715.1) under the control of the *TP53* promoter. A vector containing the same *ROR2* sequences but lacking *TP53* sequences was used as control.

CRISPR-mediated knockout of *TP53* in one mesenchymal and one epithelial cell line was performed as described elsewhere^66^. In brief, hCas9 and a guide RNA for *TP53* exon 6 were transduced into the *TERT*-immortalised cell lines human foreskin fibroblast BJ-5ta and retinal pigment epithelial cell line RPE-1 (ATCC CRL-4001, CRL-4000, LGC Standards, Middlesex, UK). The cell lines were used in the experiments immediately after purchase and were tested negative for mycoplasma. Antibiotic resistance-selected cells were single cell cloned and analysed for mutations with the Surveyor mutation detection kit (Integrated DNA Technologies, Inc., Coralville, IA). Clones with detected mutations were validated for homozygous or compound heterozygous mutations with Sanger sequencing or Nextera sequencing (Illumina). This confirmed a 19 bp deletion in *TP53* exon 6 in a BJ-5ta clone. Large genomic copy number alterations in this clone were investigated by CytoScan HD array analysis (Thermo Fisher Scientific), revealing a hemizygous deletion of distal 17p, with a break in *WRAP53*, in all cells. Thus, one *TP53* allele was deleted and the remaining allele harboured a frame-shift mutation, resulting in complete knockout of this gene. In an RPE-1 clone, two separate heterozygous mutations affecting *TP53* exon 6, where one allele harboured a 1 bp insertion and the other a 13 bp deletion, were detected. Genomic copy number analysis revealed a normal copy number of chromosome arm 17p, indicative of a complete knock-out of *TP53* via a compound heterozygous mutation. Both the BJ-5ta *TP53*^-/-^ and RPE-1 *TP53^-/-^* clone were transduced with the *TP53::ROR2* and *ROR2* vectors, respectively. The osteosarcoma cell lines Saos-2 and MG-63 were acquired as they are known to harbour the *TP53* promoter fusions *TP53::SAT2* and *TP53::ADGRE1* (previously *EMR1*), respectively^5^ (ATCC HTB-85, ATCC CRL-1427).

Bj5ta *TP53^-/-^* and RPE-1 *TP53^-/-^* harbouring either *TP53::ROR2* or *ROR2* only as well as Saos-2 and MG-63 were exposed to the DNA damaging agent cisplatin at concentrations ranging from 1-5 μM. The osteosarcoma cell line OSA was used as a control for *SAT2* and *ADGRE1* expression, respectively. Cells were harvested for RNA extraction four days following cisplatin treatment. The relative expression levels of *ROR2* (Hs00896174_m1), *SAT2* (Hs00374138_g1) or *ADGRE1* (Hs00892590_m1) were investigated using RT-qPCR and TaqMan Gene Expression assays (Thermo Fisher Scientific). The *TBP* (Hs99999910_m1) gene was used as an endogenous control. Calculations were performed using the comparative *Ct* method (i.e., ΔΔ*Ct*). The experiment was performed in biological triplicates with each replicate including technical triplicates per sample. Samples were assayed on a 7500 RT-PCR system (Thermo Fisher Scientific).

BJ-5ta wild type cells, BJ-5ta hCas9 positive cells transduced with guide RNA empty vector control (gEV), BJ-5ta *TP53^-/-^* cells, and BJ-5ta *TP53^-/-^* cells harbouring *TP53::ROR2* were cultured and harvested for RNA extraction. RNA was sequenced and analysed as described above. Unsupervised correlation-based principal component analysis was performed using the Qlucore Omics Explorer.

## Statistical calculations

Statistical calculations were performed using the two-tailed Mann-Whitney *U* test.

## Data availability

Sequencing data have been deposited at the European Genome-phenome Archive (EGA) under the accession number EGAS00001003842.

## Acknowledgements

We thank The Center for Translational Genomics at Lund University for technical support. This work was supported by the Swedish Childhood Cancer Fund, the Swedish Cancer Society, the Swedish Research Council, the Faculty of Medicine at Lund University, the Åke Wiberg Foundation, the Royal Physiographic Society (Lund, Sweden), and the Crafoord Foundation. D.B. and M.K. were supported by the Swiss National Science Foundation, the Hemmi Stiftung, the Foundation of the Basel Bone Tumour Reference Centre, the Gertrude von Meissner Stiftung, the Susy-Rückert Gedächtnis-Stiftung, the Nora van Meeuwen-Häflinger Stiftung, and the Stiftung für krebskranke Kinder, Regio Basiliensis. M.K. was supported by Slovak scientific grant agencies VEGA (grant no. 1/ 0458/18), APVV (grants no. 16/0213 and 21/0448) and from the Operational Programme Integrated Infrastructure for the project: Advancing University Capacity and Competence in Research, Development and Innovation (ACCORD) (ITMS code: 313021X329), co-funded by the European Regional Development Fund (ERDF).

## Authors’ contributions

K.H.S. and K.H.N. conceived and designed the experiments. V.P.S, O.B., E.S., M.N. and D.B. contributed tumour material and clinical information. K.H.S., M.K., and L.M. performed DNA extractions and whole-genome sequencing analyses. K.H.S., V.D., J.S. and K.H.N. analysed DNA sequencing data and interpreted the results. K.H.S., V.D., M.K., M.B., T.J., D.B. and K.H.N. performed SNP array analyses and interpreted the results. K.H.S., J.N., and L.M performed RNA extractions and RNA sequencing experiments. K.H.S., V.D. and K.H.N. carried out bioinformatic analyses of RNA sequencing data and interpreted the results. H.v.d.B., D.C.J.S. and F.F. applied low-pass whole genome sequencing on single cells. K.H.S., L.C., L.M. and J.N. conducted genomic PCR, RT-PCR, Sanger sequencing and RT-qPCR experiments, designed lentiviral vectors and performed the *in vitro* experiments. K.H.S. and K.H.N. prepared the manuscript with contributions from all other authors.

