## Supplementary Figures for "Dysregulated gene expression through *TP53* promoter swapping in osteosarcoma"

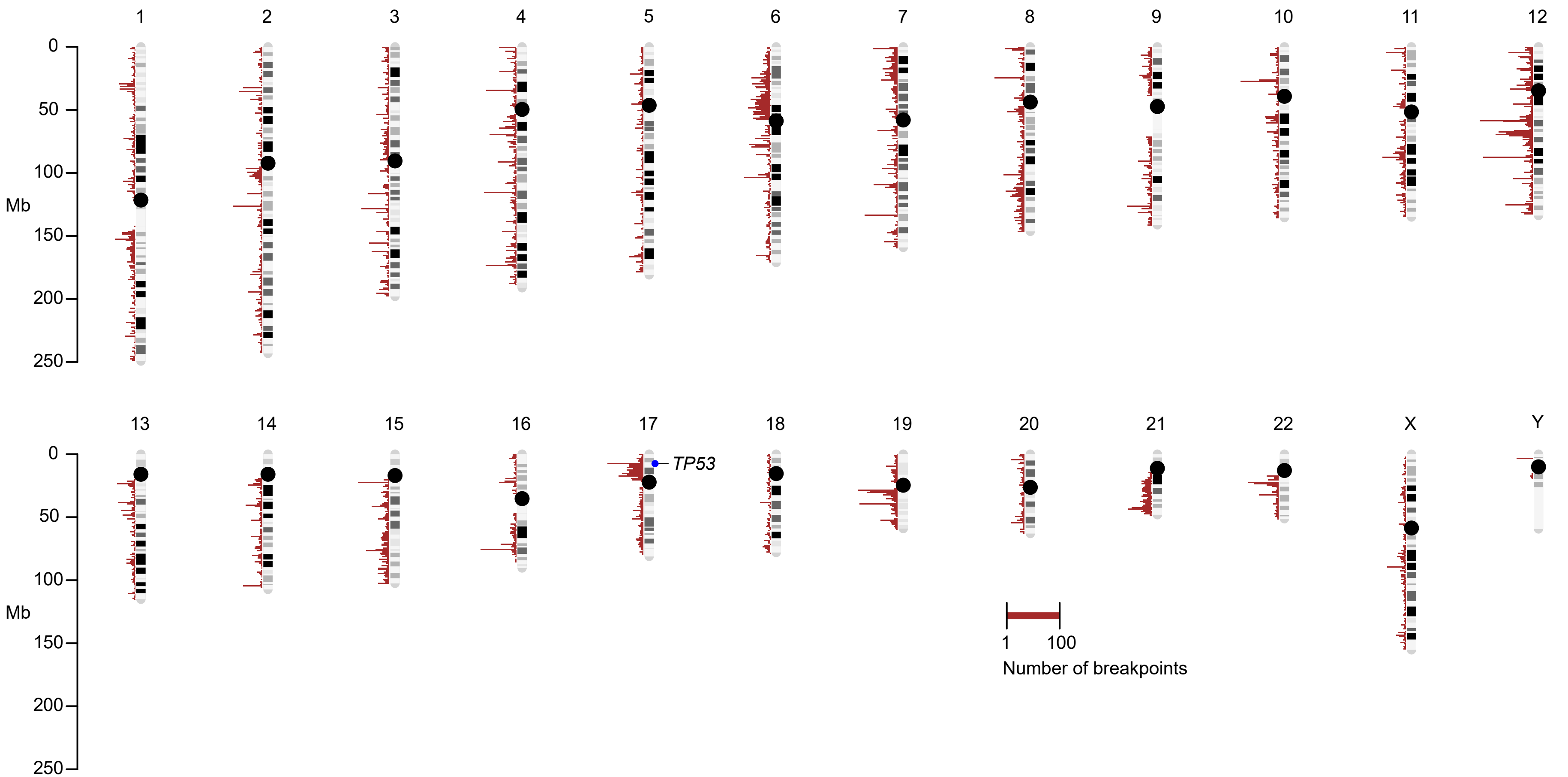


**Supplementary Figure 1. Breakpoint distribution across the 72 whole genome sequenced osteosarcomas of the discovery and validation cohorts.** Whole genome view of all detected breakpoints with a minimum support of five paired reads. Red histograms denote the amount of breakpoints detected in each genomic window. The *TP53* locus is marked on chromosome 17.

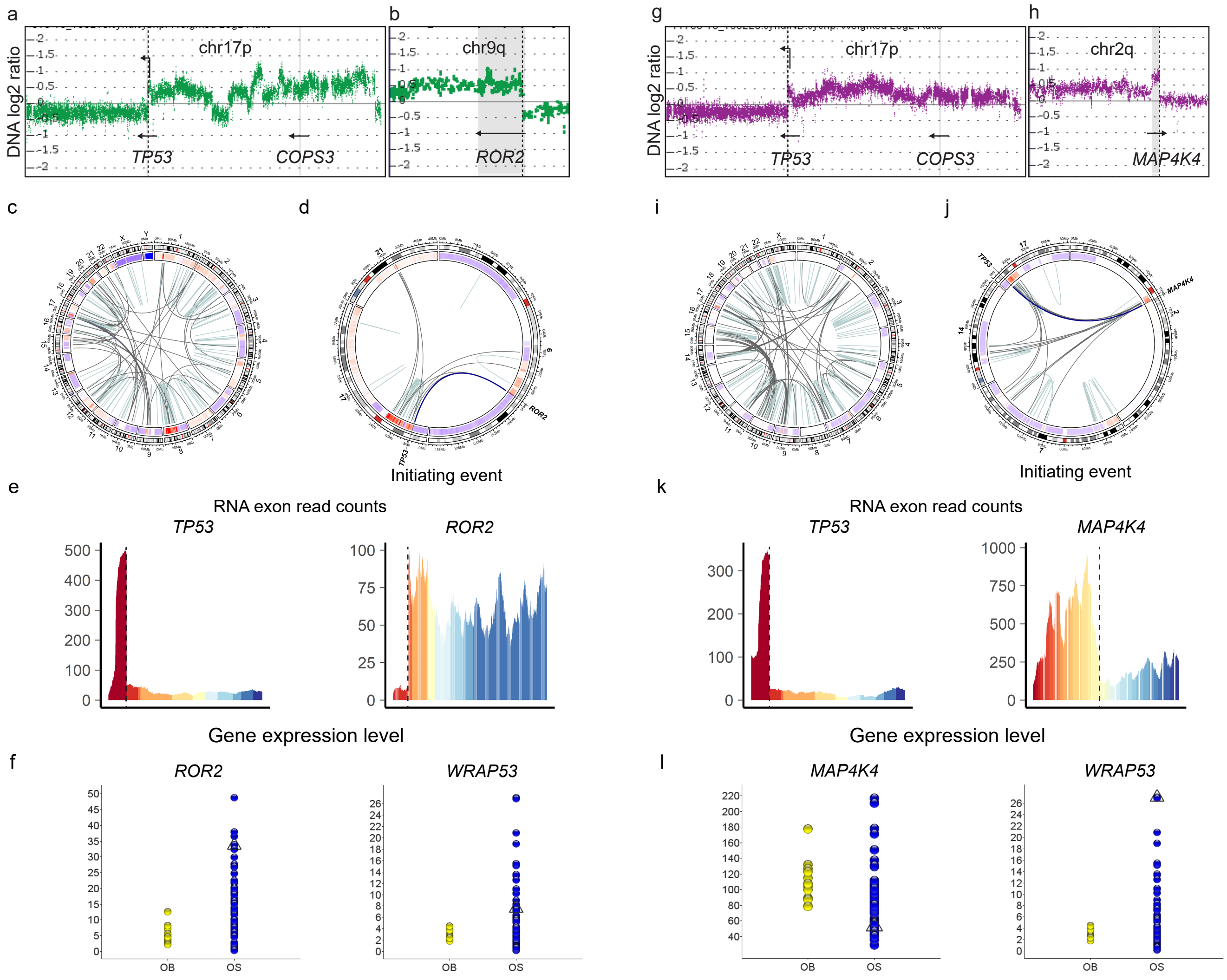


**Supplementary Figure 2. *TP53* rearrangements in selected osteosarcomas of the discovery cohort. a** Case 9, age 12. SNP array profiles of chromosome arm 17p and **b** the *TP53* promoter partner region in chromosome arm 9q, harboring the *ROR2* gene. Dotted lines indicate the fusion point. **c** Circos plots display structural variations detected by DNA mate pair sequencing in the whole genome and **d** in selected chromosomes. A blue line indicates transposition of the *TP53* promoter as the initating event that sparks a multitude of chromosomal rearrangements. **e** Exon read counts for *TP53* and the partner gene. Individual exons are plotted in different colors. Dotted lines indicate the fusion point. **f** Normalized gene expression data for *ROR2* and *WRAP53*. Case 9 is marked by a triangle. OB = osteoblastoma, OS = osteosarcoma. **g** Case 22, age 23. SNP array profiles of chromosome arm 17p and **h** the *TP53* promoter partner region in chromosome arm 2q, harboring the *MAP4K4* gene. Dotted lines indicate the fusion point. **i** Circos plots display structural variations detected by DNA mate pair sequencing in the whole genome and **j** in selected chromosomes. A blue line indicates transposition of the *TP53* promoter as the initiating event that sparks a multitude of chromosomal rearrangements. **k** Exon read counts for *TP53* and the partner gene. Individual exons are plotted in different colors. Dotted lines indicate the fusion point. **l** Normalized gene expression data for *MAP4K4* and *WRAP53*. Case 22 is marked by a triangle. OB = osteoblastoma, OS = osteosarcoma.

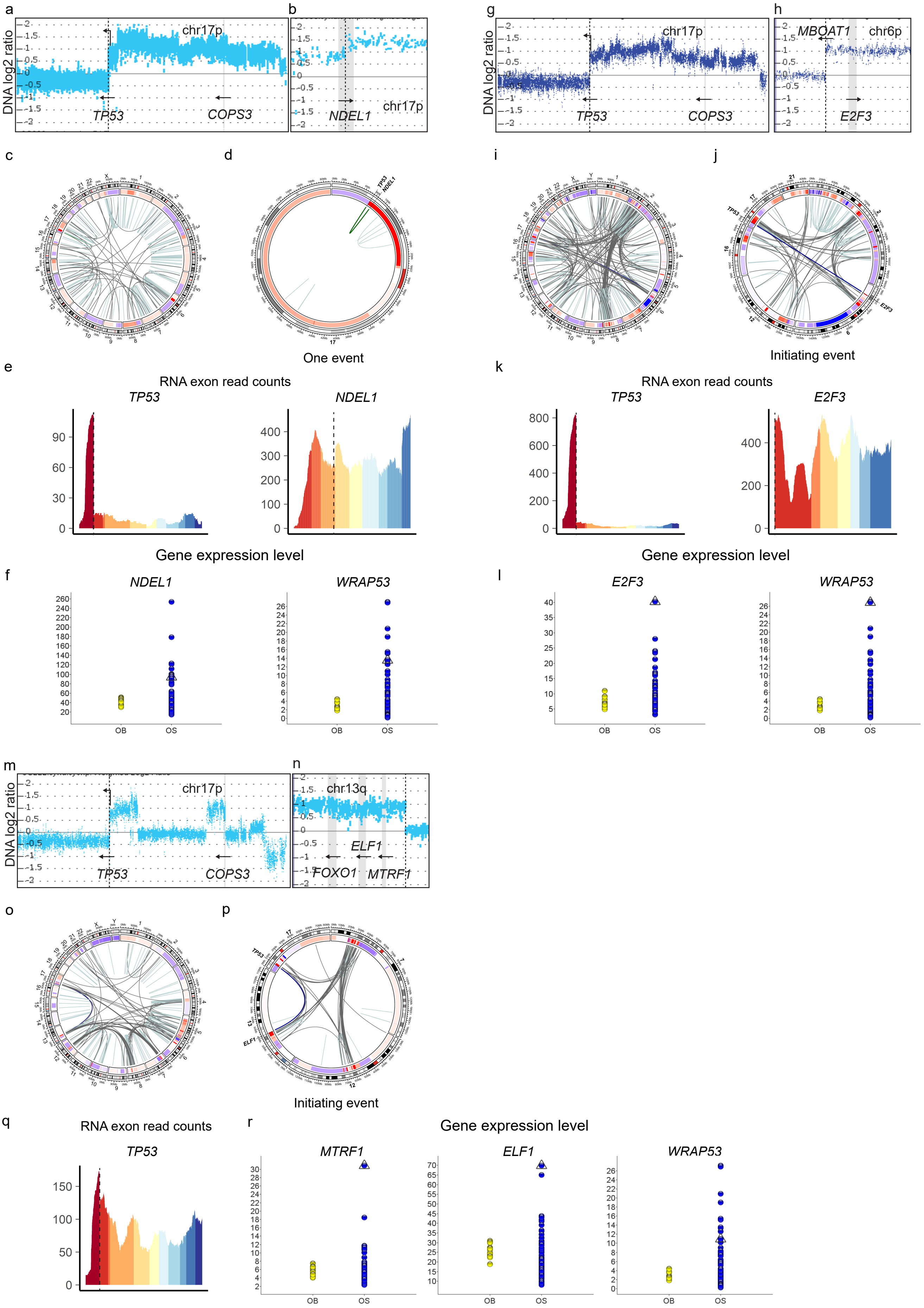
a
b
g
h
DNA log2 ratio
chr17p
TP53
COPS3
NDEL1
MBOAT1
chr6p
E2F3
c
d
i
j
One event
Initiating event
e
k
RNA exon read counts
TP53
NDEL1
E2F3
Gene expression level
f
l
WRAP53
OB
OS
m
n
chr13q
ELF1
FOXO1
MTRF1
o
p
q
r
MTRF1
ELF1

**Supplementary Figure 3. *TP53* rearrangements in selected osteosarcomas of the validation cohort. a** Case OS099, age 9, treatment-naïve diagnostic biopsy. SNP array profiles of chromosome arm 17p and **b** the *TP53* promoter partner region in chromosome arm 17p, harboring the *NDEL1* gene. Dotted lines indicate the fusion point. **c** Circos plots display structural variations detected by DNA mate pair sequencing in the whole genome and **d** in chromosome 17. A green line indicates transposition of the *TP53* promoter as a sole event. **e** Exon read counts for *TP53* and the partner gene. Individual exons are plotted in different colors. Dotted lines indicate the fusion point. **f** Normalized gene expression data for *NDEL1* and *WRAP53*. Case OS099 is marked by a triangle. OB = osteoblastoma, OS = osteosarcoma. Supplementary Figure S4 depicts *TP53* rearrangements in three lung metastases from Case OS099. **g** Case OS046, age 10. SNP array profiles of chromosome arm 17p and **h** the *TP53* promoter partner region in chromosome arm 6p, harboring the *E2F3* gene. Dotted lines indicate the fusion point. **i** Circos plots display structural variations detected by DNA mate pair sequencing in the whole genome and **j** in selected chromosomes. A blue line indicates transposition of the *TP53* promoter as the initiating event that sparks a multitude of chromosomal rearrangements. **k** Exon read counts for *TP53* and the partner gene. Individual exons are plotted in different colors. Dotted lines indicate the fusion point. **l** Normalized gene expression data for *E2F3* and *WRAP53*. Case OS046 is marked by a triangle. OB = osteoblastoma, OS = osteosarcoma. **m** Case OS222, age 17. SNP array profiles of chromosome arm 17p and **n** the TP53 promoter partner region in chromosome arm 13q, harboring the *MTRF1, ELF1* and *FOXO1* genes. Dotted lines indicate the fusion point. **o** Circos plots display structural variations detected by DNA mate pair sequencing in the whole genome and **p** in selected chromosomes. A blue line indicates transposition of the *TP53* promoter as the initiating event that sparks a multitude of chromosomal rearrangements. **q** Exon read counts for *TP53*. Individual exons are plotted in different colors. A dotted line indicates the break point. **r** Normalized gene expression data for *MTRF1, ELF1* and *WRAP53*. Case OS222 is marked by a triangle. OB = osteoblastoma, OS = osteosarcoma.

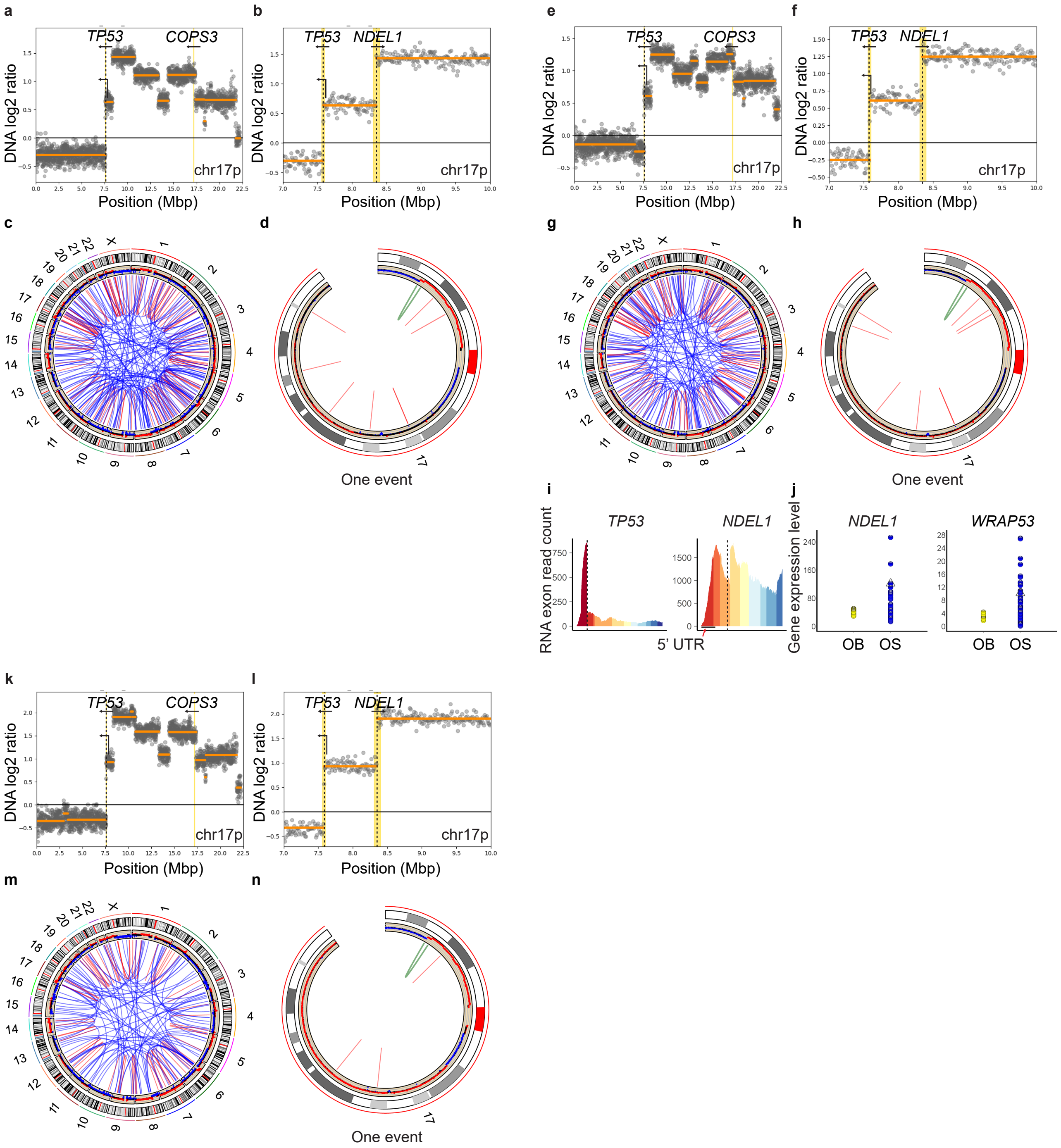


**Supplementary Figure 4. *TP53::NDEL1* in multi-sampled osteosarcoma, Case P9 (OS099). a** Lung metastasis 1. DNA paired-end sequencing of chromosome arm 17p and **b** a zoom-in on the *TP53* promoter partner region in chromosome arm 17p, harboring the *NDEL1* gene. Dotted lines indicate the fusion point. **c** Circos plots display structural variations detected by DNA paired-end sequenc-ing in the whole genome and **d** in chromosome 17. A green line indicates transposition of the *TP53* promoter as a sole event. **e** Lung metastasis 2. DNA paired-end sequencing of chromosome arm 17p and **f** a zoom-in on the *TP53* promoter partner region in chromosome arm 17p, harboring the *NDEL1* gene. Dotted lines indicate the fusion point. **g** Circos plots display structural variations detected by DNA paired-end sequencing in the whole genome and **h** in chromosome 17. A green line indicates transposition of the *TP53* promoter as a sole event. **i** Exon read counts for *TP53* and the partner gene. Individual exons are plotted in different colors. Dotted lines indicate the fusion point. **j** Normalized gene expression data for *NDEL1* and *WRAP53*. Case P9 is marked by a triangle. OB = osteoblastoma, OS = osteosarcoma. **k** Lung metastasis 3. DNA paired-end sequencing of chromosome arm 17p and **l** a zoom-in on the *TP53* promoter partner region in chromosome arm 17p, harboring the *NDEL1* gene. Dotted lines indicate the fusion point. **m** Circos plots display structural variations detected by DNA paired-end sequencing in the whole genome and **n** in chromosome 17. A green line indicates transposition of the *TP53* promoter as a sole event. Supplementary Figure 3a-f depicts *TP53* rearrangements in the treatment-naïve diagnostic biopsy from Case OS099 (P9).

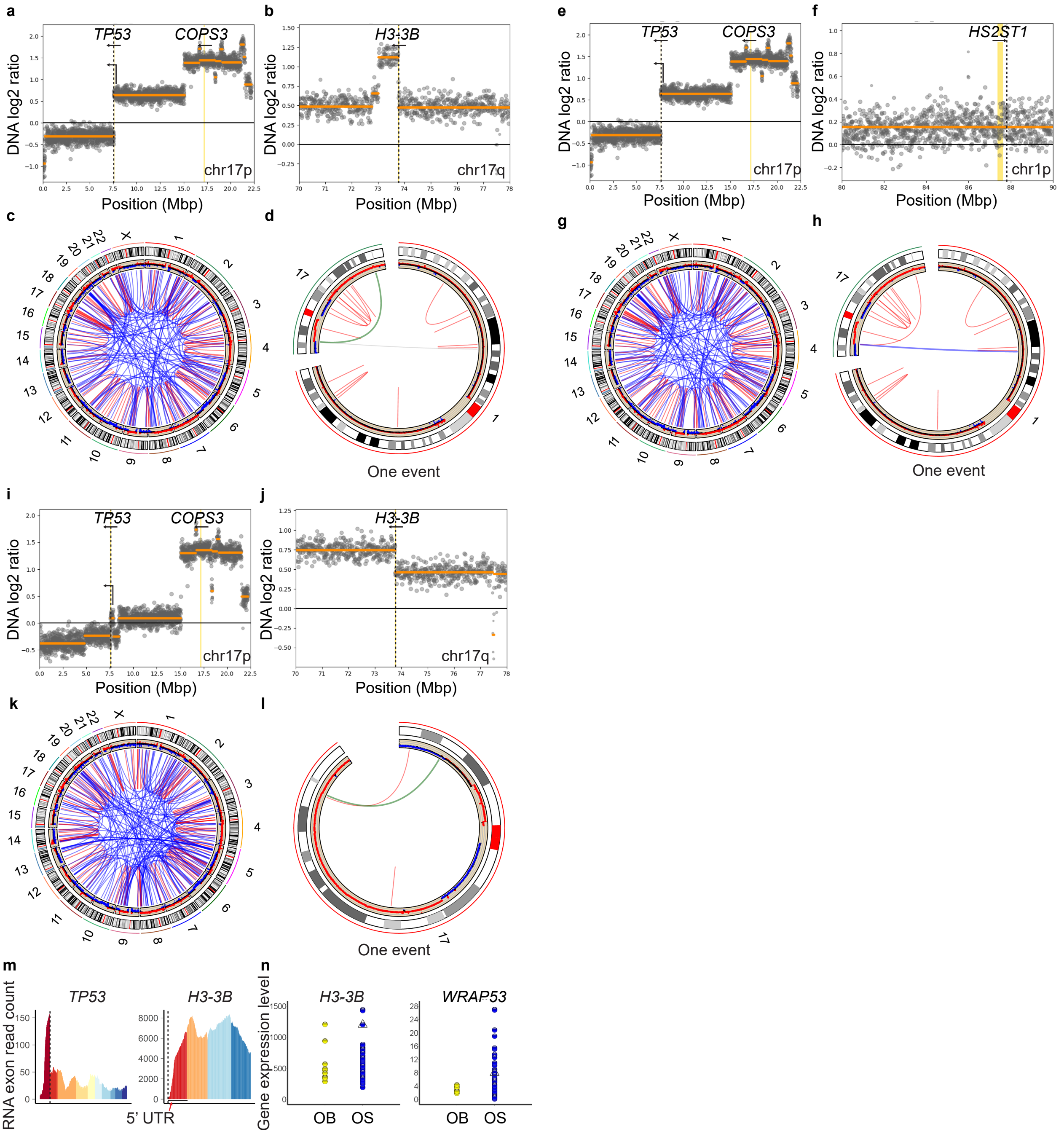


**Supplementary Figure 5. *TP53::H3-3B* in multi-sampled osteosarcoma, Case P10. a** Treatment-naïve diagnostic biopsy. DNA paired-end sequencing of chromosome arm 17p and **b** the *TP53* promoter partner region in chromosome arm 17q, harboring the *H3F3B* gene. Dotted lines indicate the fusion point. **c** Circos plots display structural variations detected by DNA paired-end sequencing in the whole genome and **d** in selected chromosomes. A green line indicates transposition of the *TP53* promoter as a sole event. **e** Treatment-naïve diagnostic biopsy. DNA paired-end sequencing of chromosome arm 17p and **f** the *TP53* promoter partner region in chromosome arm 1p, harboring the *HS2ST1* gene in opposite direction. Dotted lines indicate the fusion point. **g** Circos plots display structural variations detected by DNA paired-end sequencing in the whole genome and **h** in selected chromosomes. A blue line indicates transposition of the *TP53* promoter as a sole event. **i** Chemotherapy-treated resec-tion specimen. DNA paired-end sequencing of chromosome arm 17p and **j** the *TP53* promoter partner region in chromosome arm 17q, harboring the *H3-3B* gene. Dotted lines indicate the fusion point. **k** Circos plots display structural variations detected by DNA paired-end sequencing in the whole genome and **l** in chromosome 17. A green line indicates transposition of the *TP53* promoter as a sole event. mExon read counts for *TP53* and the partner gene. Individual exons are plotted in different colors. Dotted lines indicate the fusion point. **n** Normalized gene expression data for *H3-3B* and *WRAP53*. Case P10 is marked by a triangle. OB = osteoblastoma, OS = osteosarcoma.

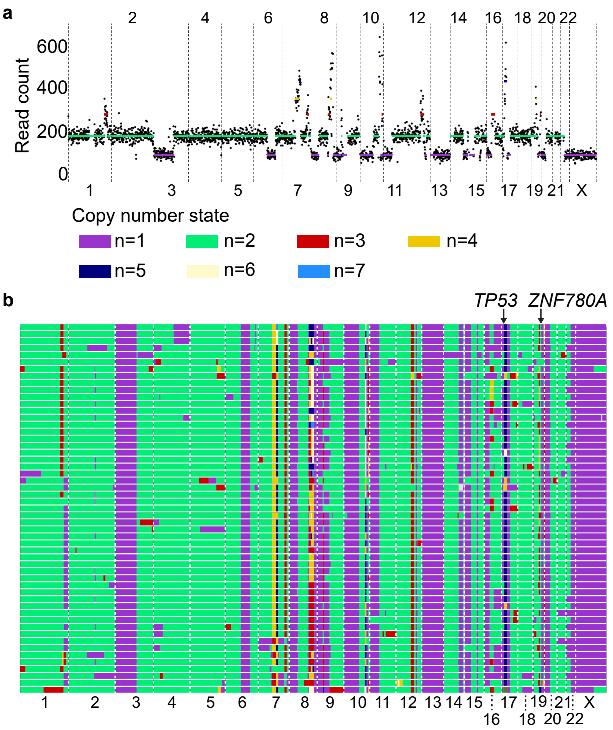


**Supplementary Figure 6. Genomic copy numbers in individual cells from a *TP53* fusion positive osteosarcoma. a** Genomic copy numbers in a representative individual cell from Case 12 harboring *TP53::ZNF780A*. **b** Heat map of genomic copy numbers across all 53 sequenced individual tumor cells. Each row of copy number states represents a single cell.

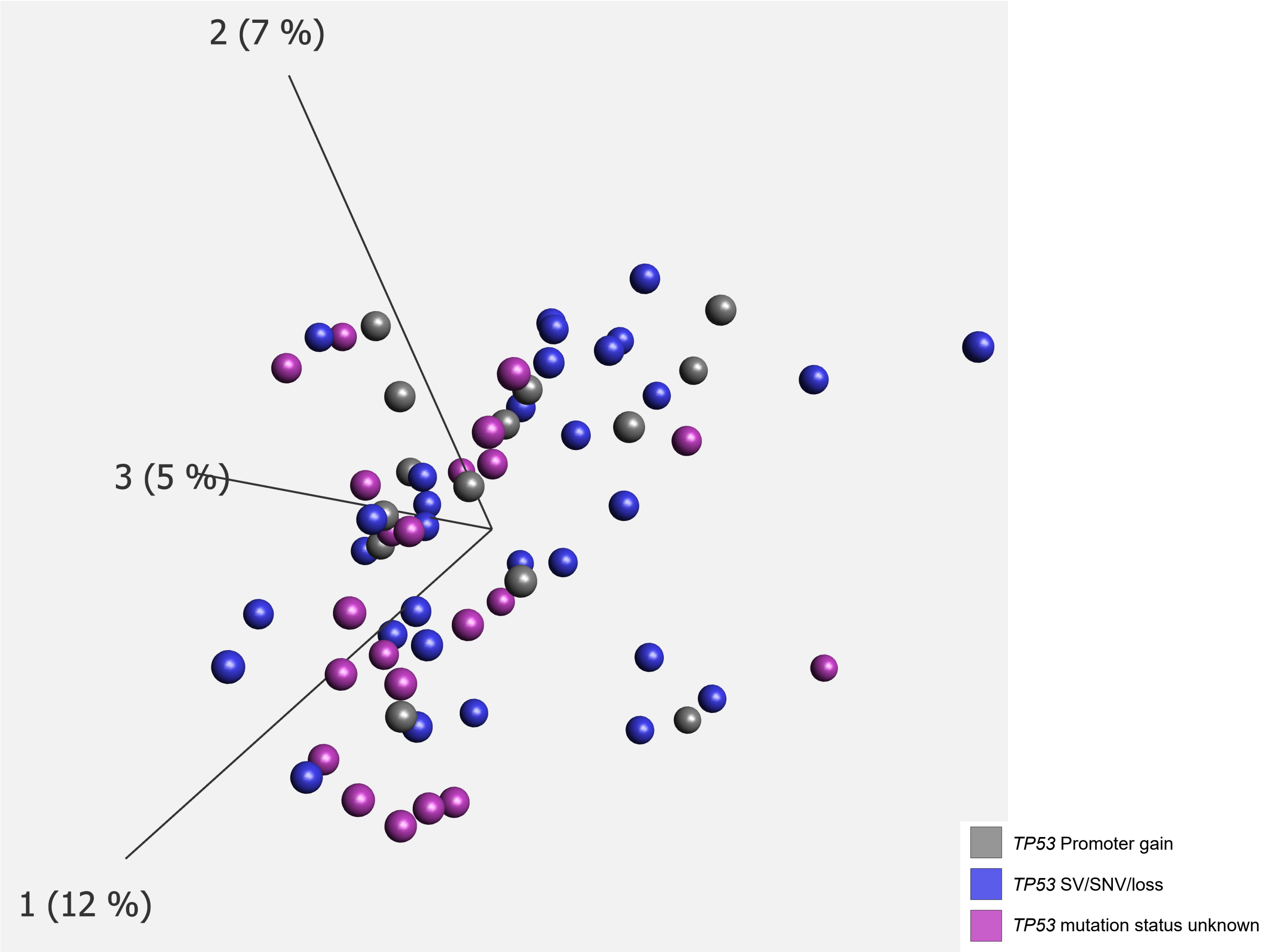


**Supplementary Figure 7. Global gene expression analyses of conventional osteosarcoma biopsies.** Unsupervised principal component analysis based on global gene expression levels in conventional osteosarcomas. Cases with *TP53* promoter gain are marked in grey. Cases without *TP53* promoter gain but with another type of *TP53* structural variant, single nucleotide variant or homozygous loss of *TP53* are marked in blue. Cases in which *TP53* status could not be fully determined are marked in pink. The first three principal components are plotted. Percentages within parentheses indicate the degree of variation represented by each axis.

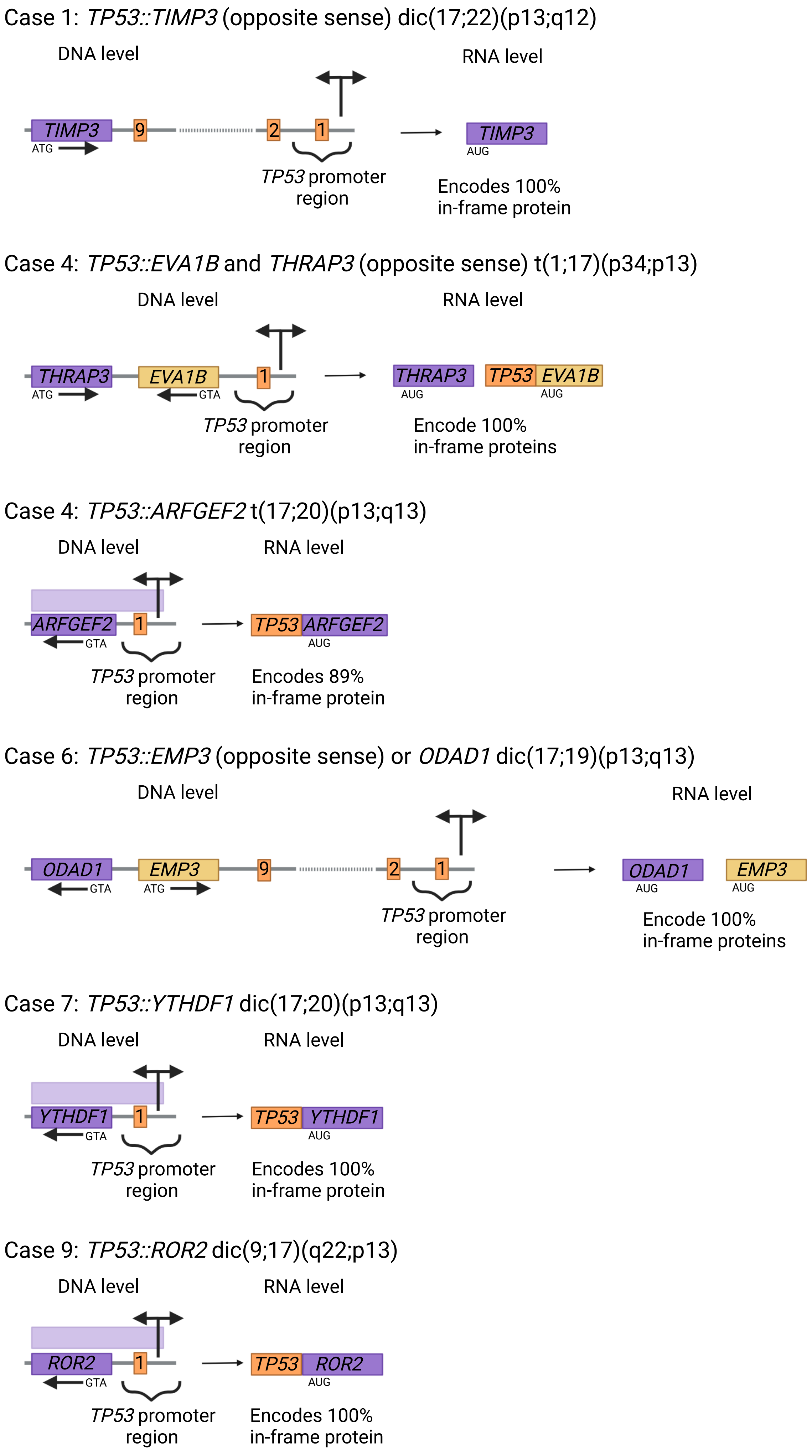
Case 1: TP53::TIMP3 (opposite sense) dic(17;22)(p13;q12)
DNA level
RNA level
TIMP3
ATG
9
2
1
TP53 promoter region
TIMP3
AUG
Encodes 100% in-frame protein
Case 4: TP53::EVA1B and THRAP3 (opposite sense) t(1;17)(p34;p13)
DNA level
RNA level
THRAP3
ATG
EVA1B
GTA
1
TP53 promoter region
THRAP3
AUG
TP53
EVA1B
AUG
Encode 100% in-frame proteins
Case 4: TP53::ARFGEF2 t(17;20)(p13;q13)
DNA level
RNA level
ARFGEF2
GTA
1
TP53 promoter region
TP53
ARFGEF2
AUG
Encodes 89% in-frame protein
Case 6: TP53::EMP3 (opposite sense) or ODAD1 dic(17;19)(p13;q13)
DNA level
RNA level
ODAD1
GTA
EMP3
ATG
9
2
1
TP53 promoter region
ODAD1
AUG
EMP3
AUG
Encode 100% in-frame proteins
Case 7: TP53::YTHDF1 dic(17;20)(p13;q13)
DNA level
RNA level
YTHDF1
GTA
1
TP53 promoter region
TP53
YTHDF1
AUG
Encodes 100% in-frame protein
Case 9: TP53::ROR2 dic(9;17)(q22;p13)
DNA level
RNA level
ROR2
GTA
1
TP53 promoter region
TP53
ROR2
AUG
Encodes 100% in-frame protein

Case 12: *TP53::ZNF780A* dic(17;19)(p13;q13)

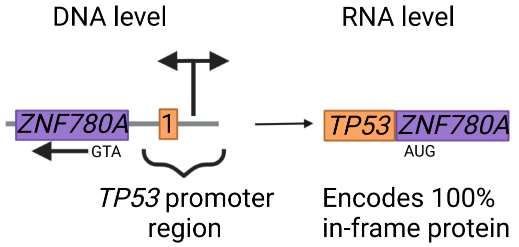


Case 13: *TP53::TGDS* or *GPC6* (opposite sense) dic(13;17)(q32;p13)

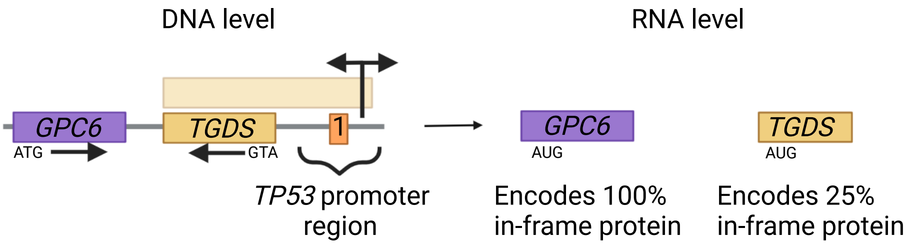


Case 17: *TP53::SUZ12* inv(17)(p13q11)

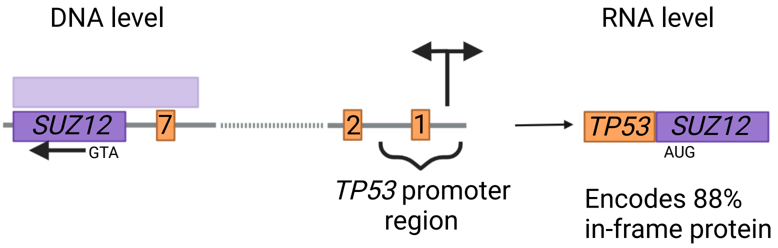


Case 22: *TP53::MAP4K4 (opposite sense)* dic(2;17)(q11;p13)

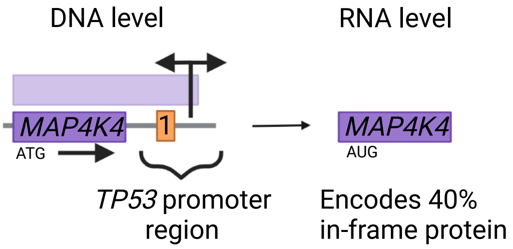


Case 24: *TP53::DAPK2* (opposite sense) or *SNX1* t(15;17)(q22;p13)

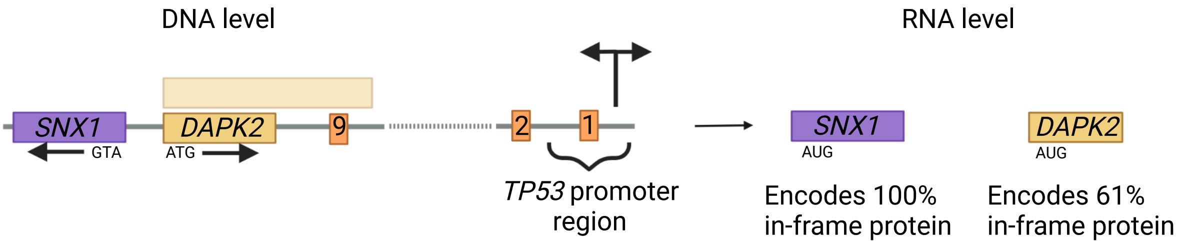

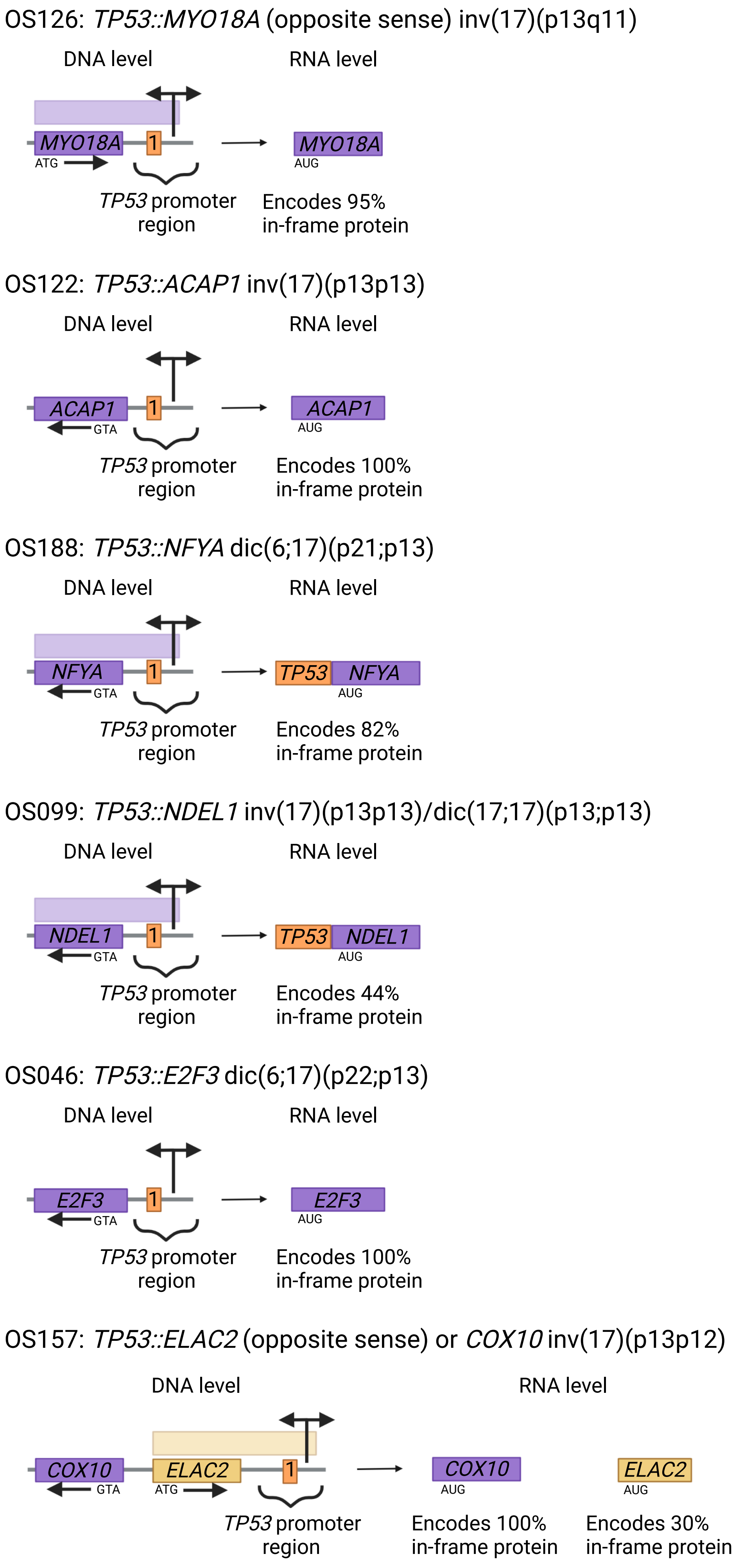
OS126: TP53::MYO18A (opposite sense) inv(17)(p13q11)
DNA level
RNA level
MYO18A
1
ATG
TP53 promoter region
MYO18A
AUG
Encodes 95% in-frame protein
OS122: TP53::ACAP1 inv(17)(p13p13)
DNA level
RNA level
ACAP1
1
GTA
TP53 promoter region
ACAP1
AUG
Encodes 100% in-frame protein
OS188: TP53::NFYA dic(6;17)(p21;p13)
DNA level
RNA level
NFYA
1
GTA
TP53 promoter region
TP53
NFYA
AUG
Encodes 82% in-frame protein
OS099: TP53::NDEL1 inv(17)(p13p13)/dic(17;17)(p13;p13)
DNA level
RNA level
NDEL1
1
GTA
TP53 promoter region
TP53
NDEL1
AUG
Encodes 44% in-frame protein
OS046: TP53::E2F3 dic(6;17)(p22;p13)
DNA level
RNA level
E2F3
1
GTA
TP53 promoter region
E2F3
AUG
Encodes 100% in-frame protein
OS157: TP53::ELAC2 (opposite sense) or COX10 inv(17)(p13p12)
DNA level
RNA level
COX10
GTA
ELAC2
ATG
1
TP53 promoter region
COX10
AUG
Encodes 100% in-frame protein
ELAC2
AUG
Encodes 30% in-frame protein

OS083: *TP53::DTD2* dic(14;17)(q12;p13)

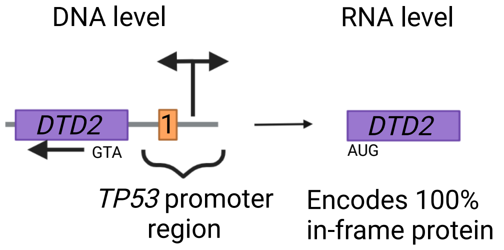


OS131: *TP53::APPL2* or *C12orf75* (opposite sense) dic(12;17)(q23;p13)

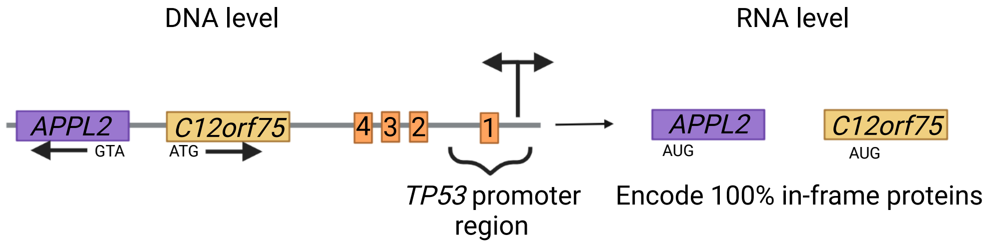


OS222: *TP53::MTRF1* or *ELF1* dic(13;17)(q14;p13)

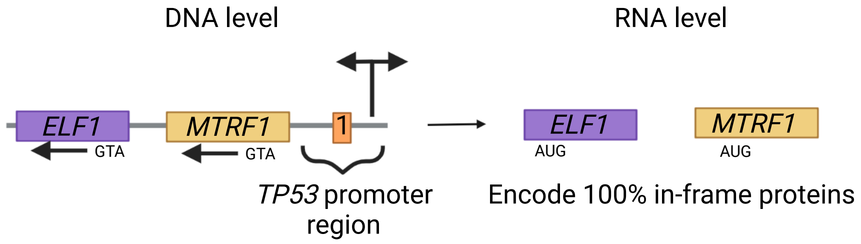


OS120: *TP53::SRSF3* or *CDKN1A* dic(6;17)(p21;p13)

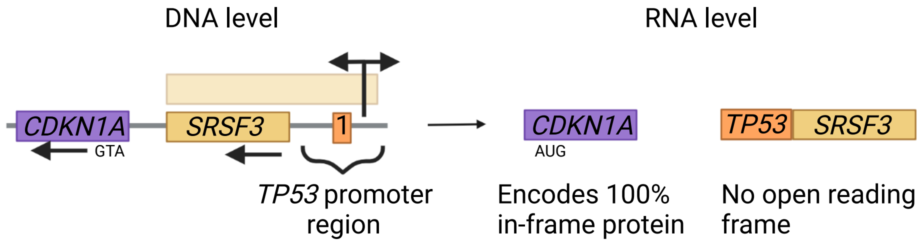


OS151: *TP53::PBX3 (opposite sense)* or *MAPKAP1* dic(9;17)(q33;p13)

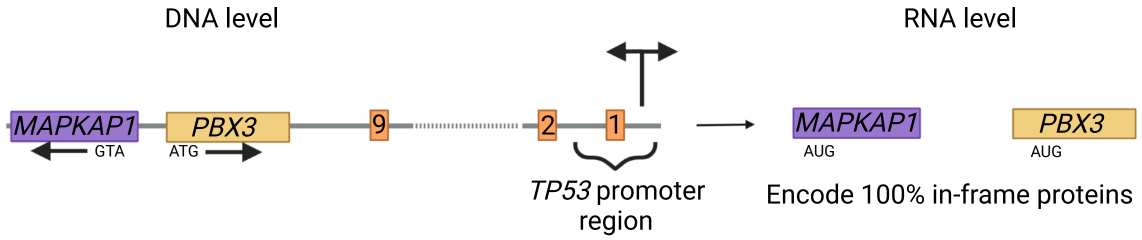


OS151: *TP53::AIPL1 (opposite sense)* or *PIMREG* inv(17)(p13p13)

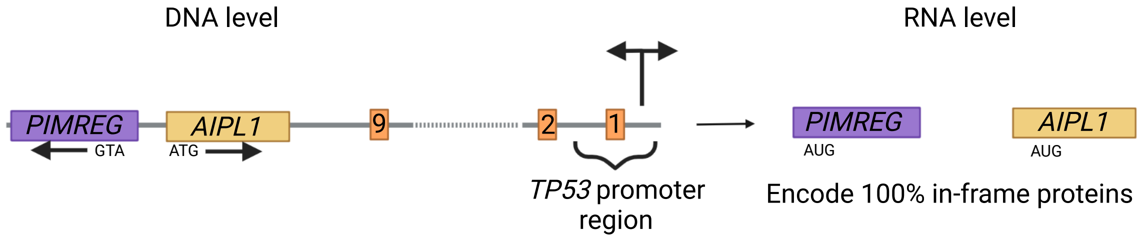

OS135: *TP53::IFT74* or *TEK* dic(9;17)(p21;p13)

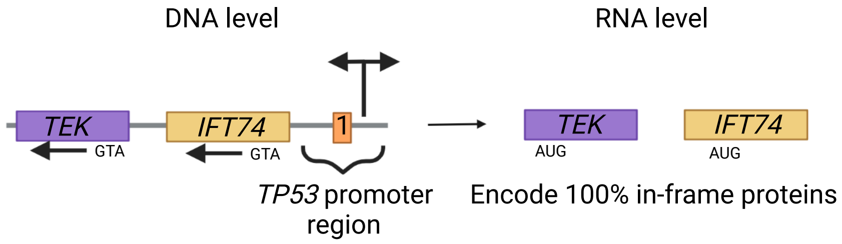


OS123: *TP53::VAMP2* or *CTC1 or PFAS* inv(17)(p13p13)

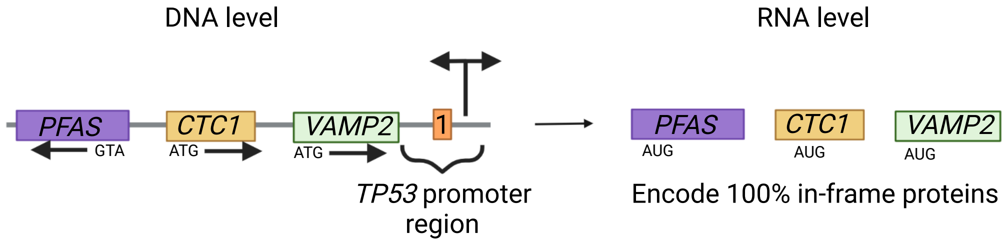


OS145: *TP53::GPC6* t(13;17)(q31;p13)

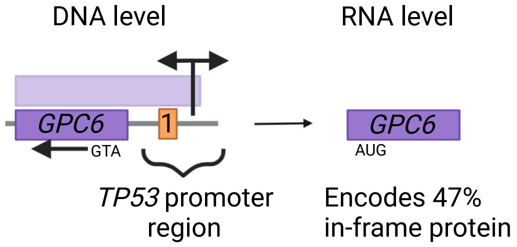


OS129: *TP53::UIMC1* or *ZNF346* (opposite sense) ins(17;5)(p13;q35q35)

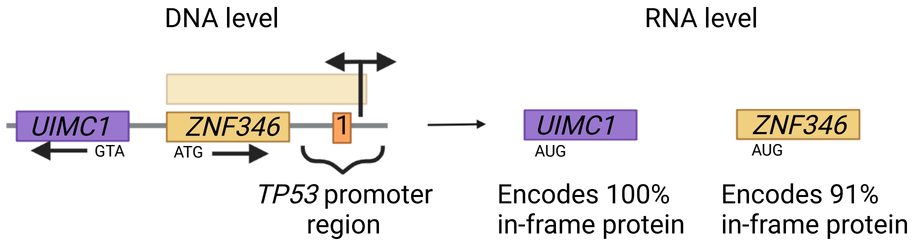


OS031: *TP53::KDM1B* or *DEK* (opposite sense) dic(6;17)(p22;p13)

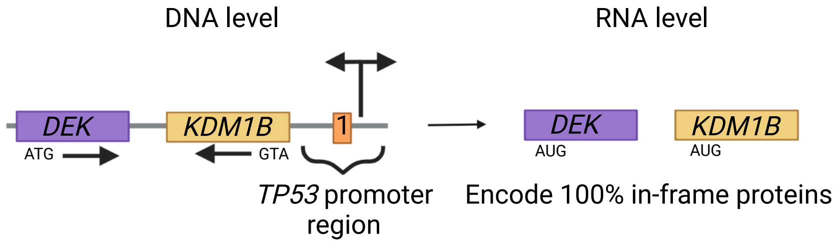


OS031: *TP53::PITRM1* t(10;17)(p15;p13)

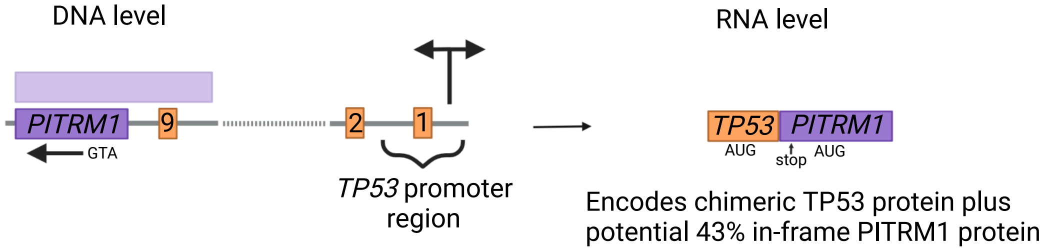

P4: *TP53::CDC5L* (opposite sense) t(6;17)(p21;p13)

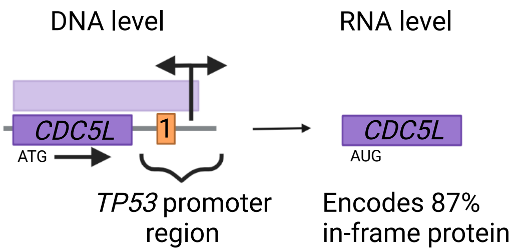


P4: *TP53::RTBDN* t(17;19)(p13;q13)

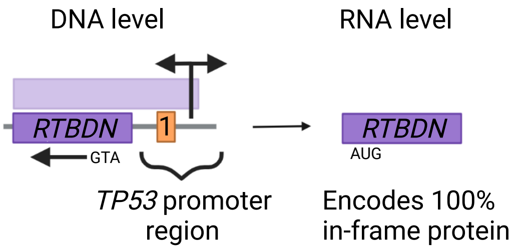


P12: *TP53::SNRPC* dic(6;17)(p21;p13)

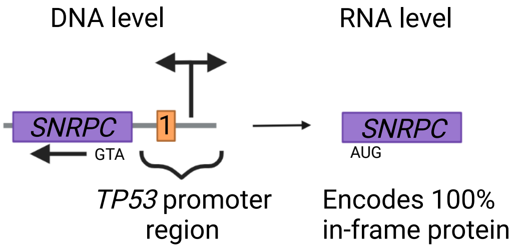


P10: *TP53::H3-3B* ins(17;17)(p13;q25q25)

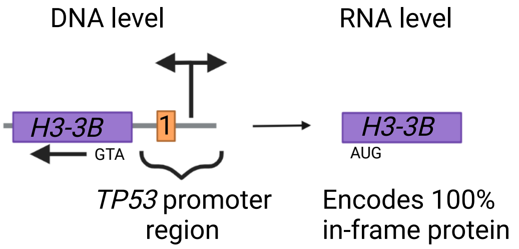


P10: *TP53::HS2ST1* (opposite sense) t(1;17)(p22;p13)

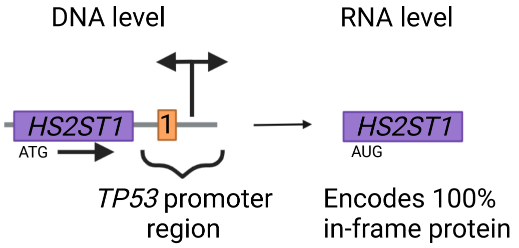


P1: *TP53::DHX8* inv(17)(p13q21)

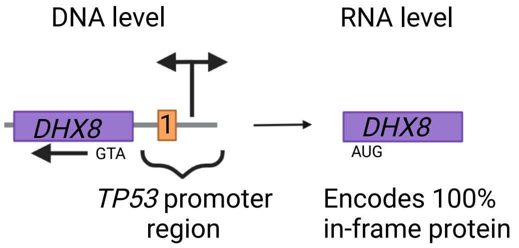

**Supplementary Figure 8. *TP53* promoter region partners in all cases where at least one partner could be identified.** The *TP53* promoter region is represented, together with the parts of the *TP53* gene involved in the structural variant according to mate pair whole genome sequencing. *TP53* exons are displayed in small, numbered orange boxes. The partner gene(s) and its/their sense are displayed to the left of *TP53*. Transparent purple boxes indicate that the breakpoint on the DNA level is within the partner gene, rather than upstream of it. The resulting mRNA molecule(s) including the approximate location of the start codon and whether it encodes a whole or partial in-frame protein are displayed to the right. In case a chimeric mRNA molecule is detected, both *TP53* and the partner gene are displayed attached to each other. Created with BioRender.com.

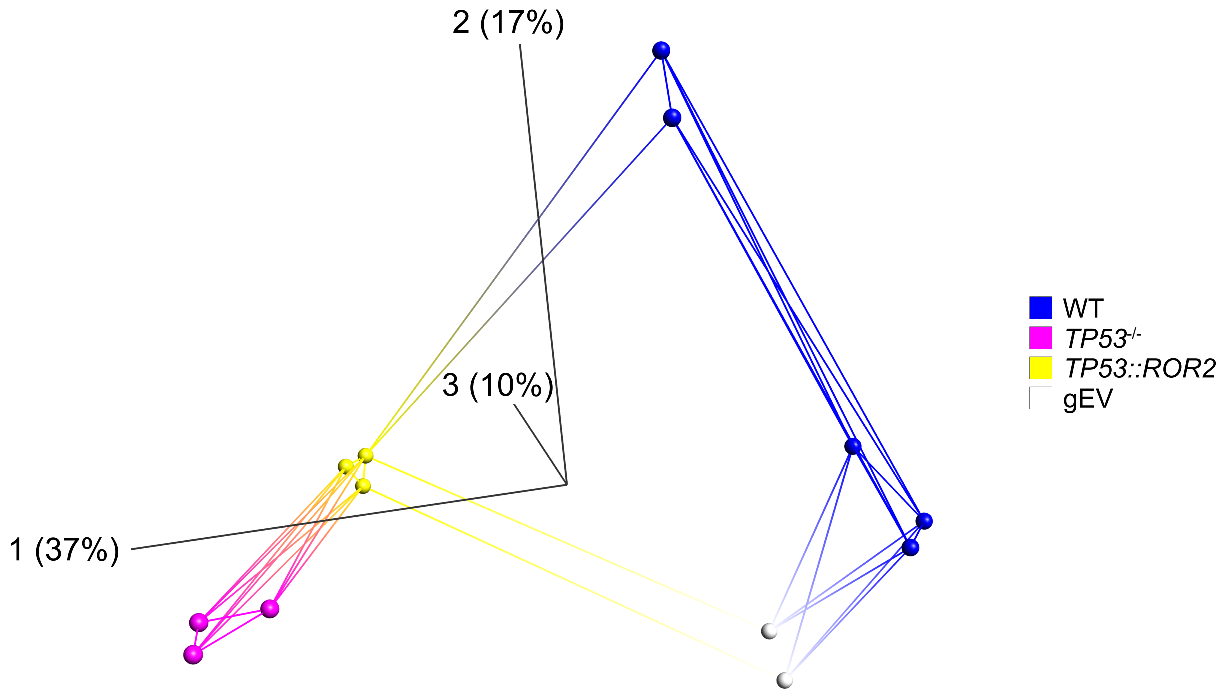


**Supplementary Figure 9. Global gene expression analyses of a *TP53*$^{-/-}$ cell model system.** Unsupervised principal component analysis based on global gene expression levels in BJ-5ta cells. Each sample is connected with its five nearest neighbors. WT = BJ-5ta wild type cells, gEV = BJ-5ta cells harboring 'guide RNA empty vector', *TP53*$^{-/-}$ = *TP53*$^{-/-}$ BJ-5ta cells, *TP53::ROR2* = *TP53*$^{-/-}$ BJ-5ta cells harboring *TP53::ROR2.* The first three principal components are plotted. Percentages within parentheses indicate the degree of variation represented by each axis.

#### Supplementary Methods

### NAFuse

NAFuse is an integrated methodology that combines and compares the output of a gene fusion detector with that of the BAM-file generated by whole genome sequencing. This allows for the detection of breakpoints in both partner genes/regions on both the RNA and DNA levels without having to manually search output files generated during downstream processing of BAM-files. The main purpose of this method is to help identify, among the huge amount of gene fusions detected with RNA sequencing, gene fusions with DNA-level support. (All the scripts and example files can be found at the following GitHub page: ***https://github.com/ValeriaDifilippo/NAFuse).***

##### Setting up NAFuse

Download BEDTools Suite (v. 2.26.0 or later) and BEDOPS [1] tools (v. 2.4.32 or later).

Based on the genome reference, download the matching gene annotation file with gene coordinates and save it in BED format. Having the reference in BED format is essential to run the bedmap command.

##### Usage

To run NAFuse apply the following steps:

**1. Prepare the genomic file.**

a. Convert sequence alignments from the DNA sequencing BAM file into BED format with the sub-commands bamTobed.
b. Annotate the BED file with the sub-command bedmap from the BEDOPS suite.
c. Split the file in two, based on the ending of the read names (/1 and /2).

Make sure that both the genomic and the gene reference file are in BED format.

The bed file generated in point *1a* contains rows with read information. It is important to notice that, when the mate information is available, each read name ends with "/1" or "/2" (*readname/1* or *readname*/2) and that they identify the two ends of one read. The two ends can align in the same gene (Supplementary Fig. 10a), in two different genes but on the same chromosome (Supplementary Fig. 10b) and in two different genes and chromosomes (Supplementary Fig. 10c).

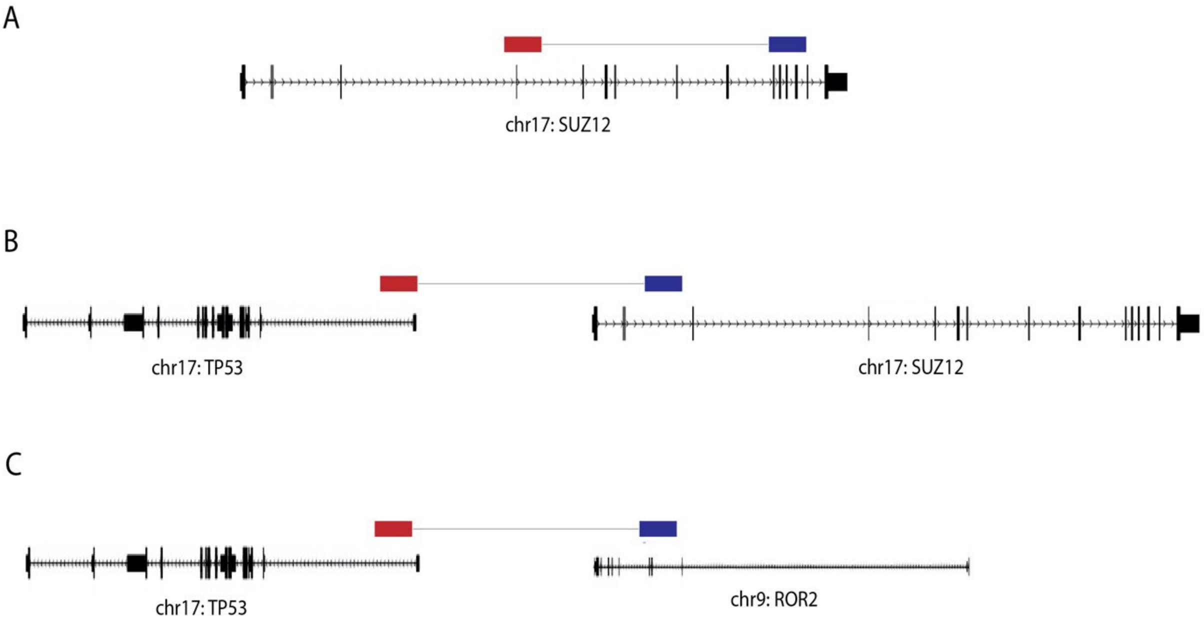


**Supplementary Figure 10. Possible ways that reads can align across the genome.**

**2. Prepare the RNA-sequencing file.**

Convert the output file from a gene fusion detector into a text file with two columns, with column one containing the 5' partner and the second column containing the 3' partner. Each line now represents a gene fusion.

**3. Now that everything is ready, it's time to run the core of NAFuse (the following script is written in Python. Example *NAFuse.py*). The script operates in two main steps.**

a. Search if there are matches between the output file from a gene fusion detector generated in point *2* and the genomic files generated in point 1c.
b. Create a file where each row contains the information belonging to the same read. It is possible to merge the lines based on read name. If a gene fusion is present in a line, it will be considered a potential gene fusion.

**4. You can now analyze you files and find new gene fusions.**

**Validation and Tips**

- NAFuse was tested in cases reported in this study. It was able to successfully detect all the gene fusions described, given that they were present in the output of a fusion detector. Gene fusion were validated by manual inspection, and some even with genomic or RT-PCR.
- In some instances, such as in Case 17 harbouring a *TP53::EVA1B* fusion, the breakpoint on the DNA level was located upstream of the 3' partner gene. To solve this, we increased gene boundaries (start and stop) by 50 000 bp each.

- Use the correct gene annotation file. Be sure to use the same gene annotation used by your gene fusion detector of choice. It can be necessary to merge multiple gene annotations files (GRCh37/hg19 and GRCh38/hg38).
